# Case-control analysis of single-cell RNA-seq studies

**DOI:** 10.1101/2022.03.15.484475

**Authors:** Viktor Petukhov, Anna Igolkina, Rasmus Rydbirk, Shenglin Mei, Lars Christoffersen, Konstantin Khodosevich, Peter V. Kharchenko

**Author notes:** Correspondence should be addressed to (P.V.K.) and (K.K.). These authors contributed equally (Viktor Petukhov, Anna Igolkina).

## Abstract

Single-cell RNA-seq (scRNA-seq) assays are being increasingly utilized to investigate specific hypotheses in both basic biology and clinically-applied studies. The design of most such studies can be often reduced to a comparison between two or more groups of samples, such as disease cases and healthy controls, or treatment and placebo. Comparative analysis between groups of scRNA-seq samples brings additional statistical considerations, and currently there is a lack of tools to address this common scenario. Based on our experience with comparative designs, here we present a computational suite (*Cacoa* – <u>ca</u>se-<u>co</u>ntrol <u>a</u>nalysis) to carry out statistical tests, exploration, and visualization of scRNA-seq sample cohorts. Using multiple example datasets, we demonstrate how application of these techniques can provide additional insights, and avoid issues stemming from inter-individual variability, limited sample size, and high dimensionality of the data.

## Introduction

The initial applications of scRNA-seq have aimed to characterize the repertoires of cell types, cell states, and continuous transitions present in different tissues (Macosko et al., 2015; La Manno et al., 2016; Usoskin et al., 2015). While these important cartography efforts are continuing to gain strength (Callaway et al., 2021; Regev et al., 2017), scRNA-seq assays have become sufficiently robust and accessible to empower many groups investigating diseases or specific biological processes (Habermann et al., 2020; Velmeshev et al., 2019). In such studies, comparative designs are most common. To characterize the impact of a particular disease, for example, a study may compare a set of samples from diagnosed patients against a set of analogous samples from healthy controls (Fig. 1A-B). More complex designs incorporating multiple sample groups, such as several patient cohorts (*e.g*. post-treatment, pre-treatment, healthy), or several disorders with similarities in pathogenesis (e.g. for neurodegeneration, Alzheimer’s, Parkinson’s, Huntington’s diseases) or several types of samples (*e.g*. tissue biopsies and peripheral blood), can be typically approached as multiple contrasts between pairs of groups. A comparison between any two or more sample groups is aimed at identifying statistically significant differences. In other words, differences that are greater than those expected from the variation within either group. However, it can be challenging to tease out statistically significant differences for clinically-oriented studies, as the differences must overcome the variability between individual patients, which can be both considerable in magnitude due to heterogeneity in pathological mechanisms even within a well-characterized disorder as well as difficult to estimate from a limited number of samples.

**Figure 1.**
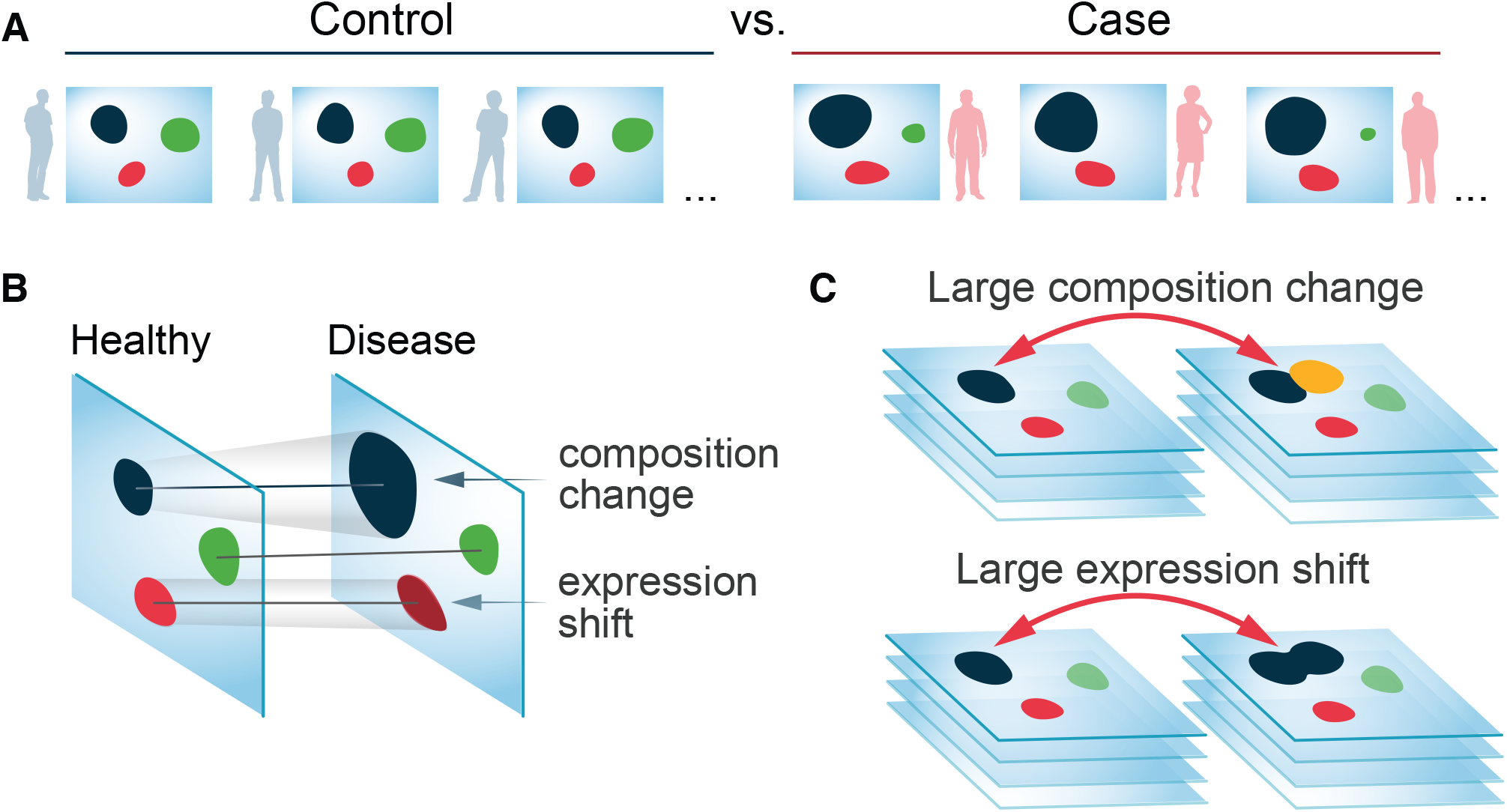
Comparative analysis of scRNA-seq collections. (A) Analysis contrasts two groups of scRNA-seq datasets, such as samples from healthy (Control) and disease (Case) individuals. (B) The two conditions may differ in state (expression shift), or proportions (compositional change) of specific cell populations. (C) Depending on the granularity of annotation, a compositional change may be captured as an expression shift.

While comparative analysis is routinely carried out using bulk RNA-seq, scRNA-seq comparisons are more challenging to interpret as they can expose multiple complementary facets on the changes occurring in a tissue (Fig. 1B). From one perspective, a contrast of two sample groups can reveal compositional differences, manifested by appearance or disappearance of cell subpopulations, or shifts in proportions of different cell types. From a different angle, the comparison can also characterize the extent and nature of transcriptional changes affecting any particular cell group. It is also possible to examine to what extent the impact of the disease is similar for different cell groups, or whether certain cell types show increased variability of their transcriptional state in disease. Here we discuss statistical considerations and potential caveats of scRNA-seq comparative analysis and propose practical methods for carrying out comparisons that we implement in an open source R package called *Cacoa* (Case-Control Analysis). We demonstrate the power of our approach by carrying out comparative analysis of five different disease-oriented datasets and revealing novel insights in pathological mechanisms that were not reported in the original studies. Though most of the results presented in the main figures are focused on the pulmonary fibrosis and multiple sclerosis datasets (Habermann et al., 2020; Schirmer et al., 2019), full results for all datasets are provided in the accompanying materials (see Datasets section of the Methods).

## Results

### Comparative analysis depends on dataset integration and clustering

The difference in the transcriptional state of cells obtained from different patients can be substantial, sometimes comparable to the difference between distinct cell types. In order to carry out comparison across conditions, it is therefore necessary to first establish correspondence between the cell populations measured in different samples. A number of computational methods have been developed to solve this “dataset alignment” problem (Butler et al., 2018; Forcato et al., 2021; Haghverdi et al., 2018), with the latest iteration of methods capable of aligning heterogeneous sample collections where not only transcriptional state, but the composition of samples may vary (Barkas et al., 2019; Korsunsky et al., 2019; Stuart et al., 2019). The alignment generated by such algorithms is used to establish a joint cell clustering, or cell annotation, so that the corresponding cell subpopulations in different samples are generally assigned to the same cluster.

Dataset alignment and clustering provide the basis for contrasting groups of samples. In general, we separate two facets of comparative analysis: i) analysis of compositional shifts, that examines changes in the frequency of different cell subpopulations, and ii) analysis of transcriptional shifts, that characterizes the changes in the state of individual subpopulations (Fig 1A-B). The two perspectives are to some extent complementary: a significant alteration of a cell type, such as B cell activation, may be viewed as a transcriptional or compositional shift, depending on whether the activated B cells are annotated as a distinct subpopulation (Fig. 1C). As granularity of cell clustering is generally determined by the investigator or the choice of the algorithm and parameters, it may obscure some of the trends in the data. In performing different comparisons between conditions, we therefore complement analysis of annotated cell clusters (cluster-based) with more general methods for performing comparisons on the dataset alignment itself (cluster-free).

### Analysis of compositional differences based on annotated clusters

The most common approach to analyze compositional differences simply compares the proportions of different cell types between conditions. For example, a recent study of pulmonary fibrosis (PF) has contrasted 20 diagnosed patients with 10 non-fibrotic controls (Habermann et al., 2020). To characterize the impact of the disease on the composition of the lung tissue, the authors have analyzed the relative proportions of different epithelial cell types. The study noted a significant decrease in the proportion of type 1 alveolar (AT1) and transitional type 2 (Transitional AT2) epithelial cells in the disease state, using Mann-Whitney U tests to evaluate statistical significance of the proportion differences. Such comparisons can reveal statistically significant differences in cell type abundance even for studies with a modest (∼10) number of samples, illustrating the striking extent of remodeling in many diseases (see Datasets section for full reports). Simple comparison of proportions, however, has several potential complications. First, the proportion tests for different cell types are not independent of each other. For example, a significant expansion of a single cell type can lead to a statistically significant decrease in proportion of all the other cell types (Fig. 2D,E).

**Figure 2.**
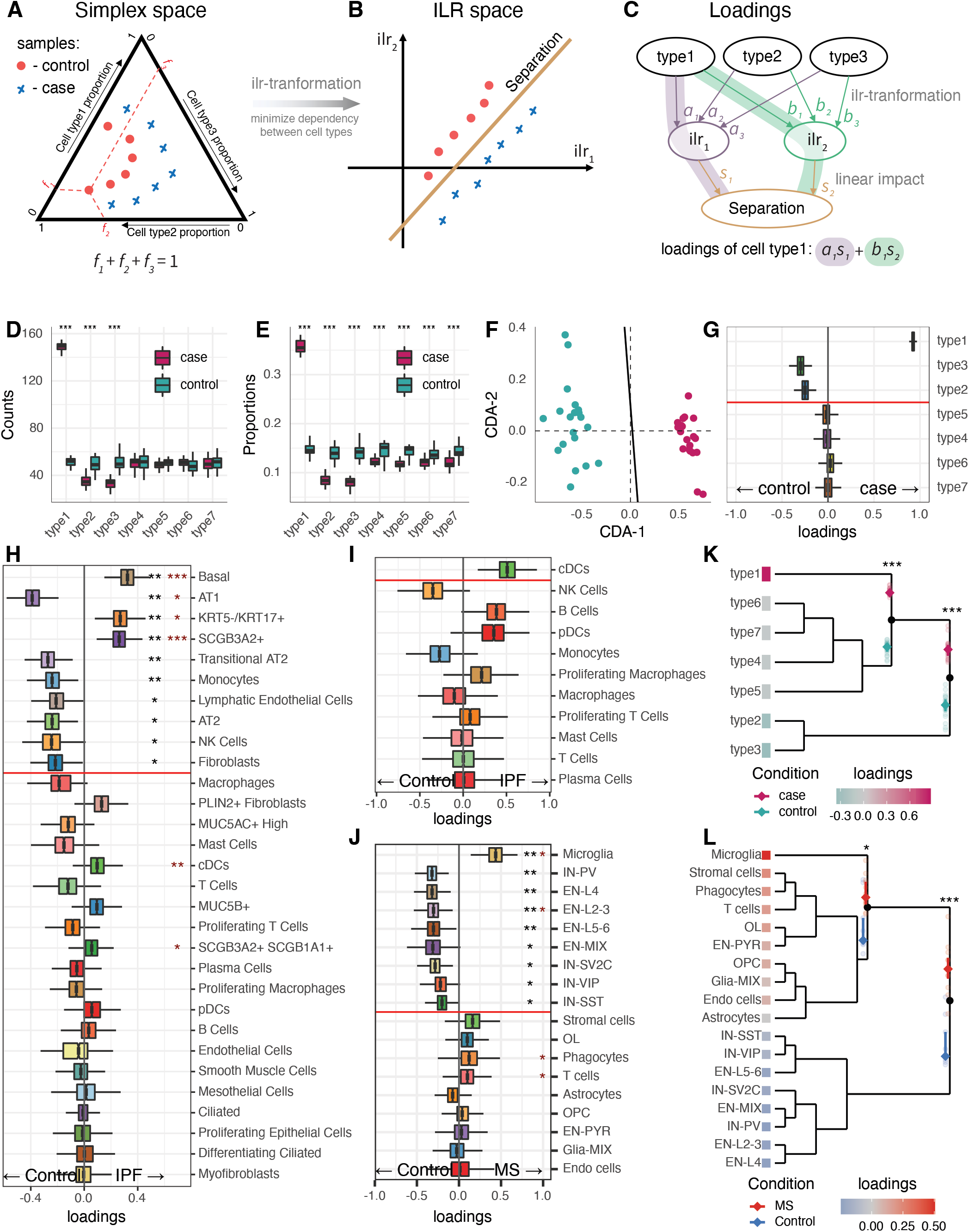
Compositional data analysis for scRNA-seq experiments. (A-C) Schematic illustration of compositional data analysis (CoDA). (A) Proportions of different cell populations form a simplex space (*i.e*., add up to 1). (B) CoDA applies ilr transformation to reduce dependencies between variables, and looks for a surface that optimally separates case and control samples within that space. (C) Since the separating surface is defined in ilr-transformed space, each cell type can contribute to the surface through different variables. Cell type loadings quantify the overall contribution of the cell type to the separation of the two conditions based on all of the transformed variables. (D-E) A toy example illustrating problems with analysis of proportions. It shows seven cell types (x-axis), the first three of which change between conditions. Absolute counts and cell type proportions are shown in (D) and (E), respectively. After the normalization all cell types show significant differences (E). (F) Visualization of CoDA separating surface, visualized on the first two principal components of the ilr-transformed space. (G) CoDA loadings (x-axis) for the cell types (y-axis) in the simulated example. Positive loadings correspond to over-representation in the Case and negative loadings mean over-representation in Control. The boxplots depict the uncertainty of the loading coefficients under cell and sample subsampling (see Methods). The red horizontal line separates cell types passing significance threshold (*p <* 0.05 after BH correction). These loadings match closely to the original change (D), with only the affected cell types passing the significance threshold. (H) CoDA loadings for the PF dataset, same representation as (G). Dark-red asterisks represent the significance level of the simple proportion test with black asterisks showing CoDA significance. All alveolar epithelial populations are significantly decreased in IPF, while simple proportion analysis detected only AT1 cell type. (I) CoDA analysis with the scope reduced only to CD45+ cell types. (J) CoDA loadings for the MS dataset. CoDA detected decrease in all neuronal populations, while only EN-L2-3 was detected by proportion analysis. (K-L) Hierarchical representation of the compositional changes on the toy example (K) and the MS dataset (L). The toy dataset groups three significantly different clades: ‘type 1’, ‘type 2’ + ‘type 3’ and the rest, which matches the simulation design. In the MS dataset, the hierarchy shows significant separation between neuronal and glial cell types. Significance codes: \**p <* 0.05, ** *<* 0.01, * * * *<* 0.001.

To overcome this limitation we followed an approach of compositional data analysis (CoDA), translating the abundances of N cell types into N-1 simplex space using isometric log-ratio transformation (*ilr*) (Pawlowsky-Glahn et al., 2015) (Fig. 2A-B). As ilr transformation uses normalization by geometric mean of abundances, the resulting N-1 variables are largely independent and thus more suitable for statistical comparisons. To compare the composition between the disease and healthy samples, we identify a surface within this N-1 simplex space that optimally separates the conditions being compared (Fig. 2B), and estimate the extent to which different cell types drive the orientation of that surface (loading coefficients, Fig. 2C). To estimate the statistical significance of loading coefficients, we generate an empirical background distribution based on random resampling of samples and cells across the entire dataset (see Methods). The resulting compositional analysis approach improves robustness and sensitivity of the compositional tests (Fig. 2D-G).

Applying this compositional analysis to the full pulmonary fibrosis (PF) dataset, we detect significant reduction in AT1 and Transitional AT2 epithelial populations noted in the original publication (Fig. 2H). We also find significant reduction in the AT2 cells, demonstrating that all alveolar epithelial populations decrease in abundance in PF. The reduction was most pronounced in the AT1 cells, consistent with AT1 being particularly sensitive to lung injury that leads to their cell death by apoptosis (Zemans et al., 2015). The strongest increase was observed for the Basal cells, whose predominance is a well-known predictor of PF (De Andrade and Luckhardt, 2017; Martinez et al., 2017). Increase in SCGB3A2+ and KRT5-/KRT17+ has been also associated with lung injury in PF and was reported in the original study (Habermann et al., 2020).

Applying Cacoa compositional analysis to another example dataset, contrasting 12 cortical samples from patients affected by multiple sclerosis (MS) with 9 controls (Fig. 2J) (Schirmer et al., 2019), we observed as in the original study, that the strongest difference was the increase of microglia cell abundance. However, we additionally found that excitatory (cortical principal) neurons across all cortical layers and some inhibitory (GABAergic) neuron types (IN-PV, IN-VIP) were significantly decreased in MS samples (Fig. 2J), while the original paper pointed to only excitatory L2-L3 neurons using a simple proportion analysis. The decrease in the inhibitory neurons in MS revealed by Cacoa analysis correlates well with recent immunohistochemical data showing a general reduction of inhibitory neurons in the cortex of MS patients, in particular IN-PV (Zoupi et al., 2021). Since IN-PV provide the strongest perisomatic inhibition to excitatory neurons, such reduction in IN-PV should disinhibit excitatory neurons, which is supported by reduction in inhibitory synapses in the cortex of MS patients (Zoupi et al., 2021).

While transformation into ilr simplex space improves robustness to outliers, the analysis of relative cell type abundances ultimately depends on the scope of populations being considered. For instance, the investigator may be interested in the compositional shifts within the immune compartment alone and perform compositional analysis limiting it to immune populations. In the PF dataset, analysis of such reduced scope reports a significant increase in the abundance of conventional dendritic cells (cDCs) cells in disease (Fig. 2I), whereas in the original full scope analysis the cDCs abundance was unchanged (Fig. 2H). Increase in cDCs is consistent with the literature showing enhanced recruitment of DCs to PF lungs due to chemokine-driven infiltration into fibrotic tissue (Marchal-Sommé et al., 2007). In general, both types of scope analysis are equally valid, as they address different questions. Technically, the biggest difference occurs due to a choice of unchanged or *reference* cell types. Cacoa compositional analysis determines reference cell types as a non-contrasting (i.e. not separating according to condition) group of cells with loading coefficients closest to zero, which given the geometric nature of the ilr space tends to select the most stable group. Alternatively, reference cell type(s) can be provided by the investigator, to carry out compositional analysis under the assumption that these cell types have not changed in their absolute abundance.

Interpreting compositional shifts in tissues with many cell types can be challenging. Some groups of cell types may change to a similar extent, which may aid the overall interpretation. To capture such congruent behavior, we derive a hierarchical representation of the observed compositional shifts, using a greedy strategy to group cell types that are affected in the same way and produce a simplified view of the shifts in the dataset. For instance, in the simulated example (Fig. 2D-E), type2 and type3 were shifted in the same manner, and would be placed in the separate clade, significantly different from the other cell types (Fig. 2K). In the MS study, this analysis reveals a parsimonious perspective on the changes, capturing a reduction in abundance of all neuronal types in one clade, and an increase in glial types in the other (Fig. 2L).

### Cluster-free compositional differences

As with all “cluster-based” analysis, the granularity of the annotated cell types limits the resolution at which the changes can be detected and interpreted. If the disease, for instance, impacts only a specific subset of an annotated cell type, this may not be evident from cluster-based analysis. Furthermore, annotation of disease samples might be hampered by a lack of knowledge in biology of the disease, which could compromise annotation. This limitation can be addressed by “cluster-free” analysis, which compares densities of cells within the expression manifold between the samples from disease and control conditions (Fig. 3A-C). In its simplest form, the expression manifold can be approximated by a two-dimensional embedding (e.g. UMAP or t-SNE). More general approximation of the manifold is provided by the graphs generated by the dataset alignment methods (Butler et al., 2018; Haghverdi et al., 2018). To evaluate compositional differences between conditions, for each sample we will estimate a normalized cell density in embedding or graph space (Fig. 3A), and then assess the magnitude and statistical significance of the sample density differences between conditions (Fig. 3B,C). While for embeddings cell density can be readily estimated using the standard kernel density estimation (KDE) procedure (Chacón and Duong, 2018), in graph space we make use of a KDE generalization, formulated as a filter in the graph Laplacian space (Burkhardt et al., 2021; Shuman et al., 2013). The statistical comparison of the densities between conditions is carried out against an empirical background distribution, generated by randomizing samples across conditions. Cluster-free analysis requires a different procedure for adjusting p-values for multiple comparison, as doing so in a naive manner would lead to nearly complete lack of sensitivity. We therefore implemented p-value correction based on the distribution of the maximal statistic under permutations of condition labels (Holmes et al., 1996) (see Methods).

**Figure 3.**
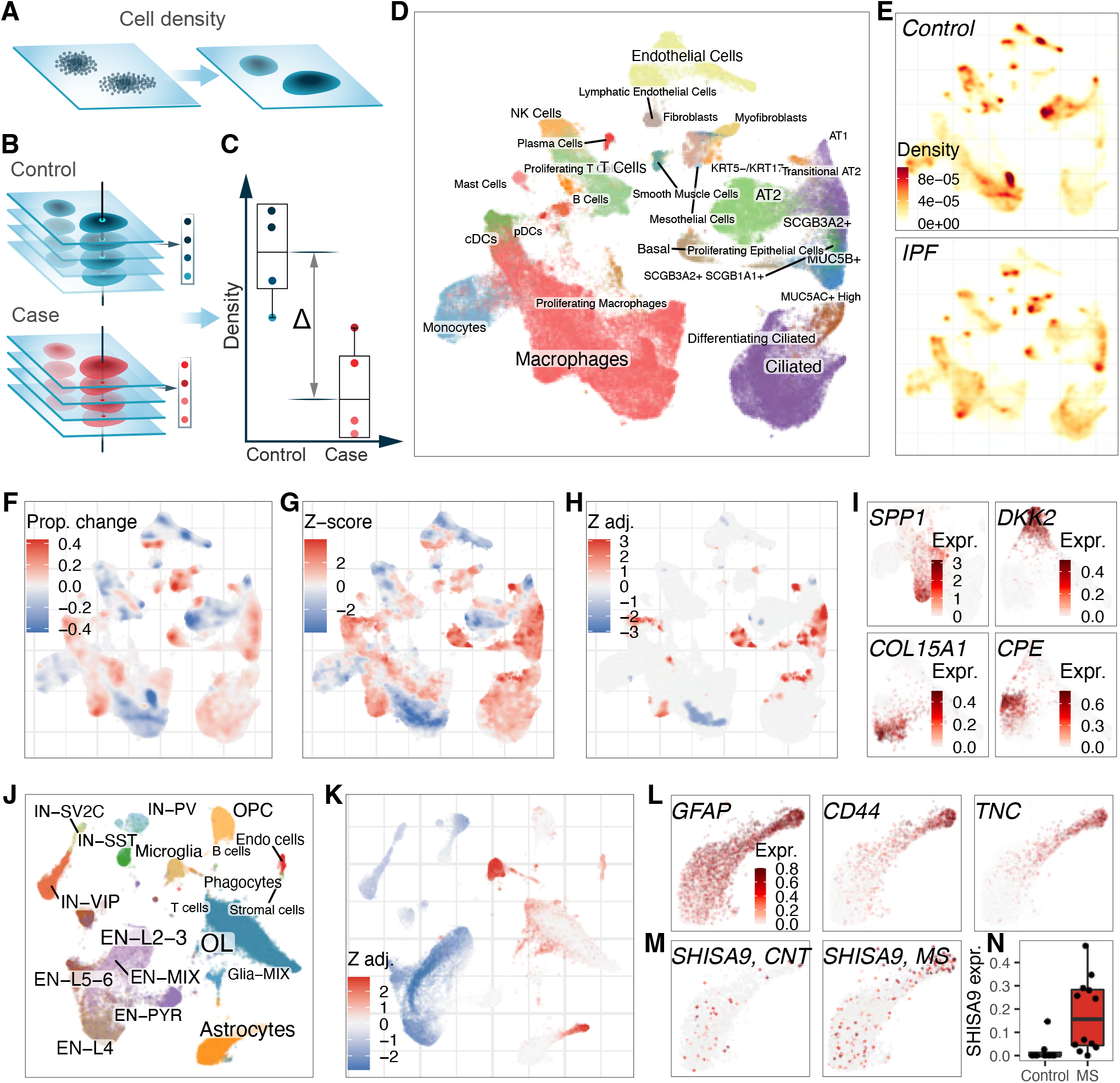
Cluster-free analysis of cell type composition. (A-C) Schematics of cluster-free compositional analysis. (A) Cell density is estimated for each sample, either in the space of a 2D embedding or on a cell graph. (B) Each point has a density value for each sample. (C) These values are then used to evaluate statistical difference between the two conditions at each point. (D) Annotated UMAP embedding for the PF dataset. (E) Visualization of the average cell density within the Control (top) and IPF (bottom) conditions, using embedding density estimation. Darker colors correspond to denser regions. (F) Magnitude of proportion change between the Control and IPF is visualized on the UMAP embedding. The color shows the average density difference, normalized by the maximum density across all points. (G) Separability z-score, (as in panel C), estimated for the PF dataset. These scores are intermediate values, used for further estimation of statistical significance (see H), and can be interpreted as significance values un-adjusted for multiple hypotheses. (H) Adjusted significance levels are shown for the PF dataset, converted into z-scores of the standard normal distri-bution for the visualization purposes. Colors show points with the absolute adjusted z-score values above 1.6 pass significance threshold of p=0.05. (I) Marker genes for Macrophage (SPP1) and Endothelial (DKK2, COL15A1 and CPE1) subpopulations. Darker colors correspond to higher expression levels. (J) Annotated UMAP embedding for the MS dataset. (K) Adjusted significance of changes is shown for the MS dataset, using graph-based density estimates. The format is similar to (H). (L) Visualization of Astrocyte marker genes for Astrocytes subpopulation. (M-N) Changes in SHISA9 gene expression between control and MS conditions. (N) shows aggregated expression (y-axis) by samples (dots) within each condition (x-axis and colors).

Cell density analysis of the PF dataset recapitulates the major shifts in abundance captured by cluster-based analysis, including marked increase of Basal, SCGB3A2+ and MUC5B epithelial populations (Fig. 3D-H). In some cell types, however, comparison between conditions reveals subpopulations with distinct behavior. For instance, while the macrophage population as a whole remains relatively unchanged, a specific subset of macrophages shows significant increase in PF relative to control samples (Fig. 3H). This subset is marked by expression of osteopontin (*SPP1*, Fig. 3I), which corresponds to pro-fibrotic macrophages known to proliferate rapidly in PF patients (Morse et al., 2019; Wynn and Vannella, 2016). Similarly, a subpopulation of endothelial cells (Fig. 3I), marked by expression of *COL15A1* - systemic-venous endothelial cells (Schupp et al., 2021), also shows an increase in PF compared to control samples, while arterial endothelial cells, marked by *DKK2* (Schupp et al., 2021), are significantly decreased in PF.

Cluster-free compositional analysis of the MS dataset also recapitulates the difference in major subpopulation abundance that was captured by the cluster-based analysis, such as overall reduction in neuronal abundance relative to glial populations (Fig. 3J,K). Importantly, cluster free analysis additionally reveals a “tail” of astrocyte population that is significantly increased in MS. This astrocyte subpopulation expresses *GFAP, CD44* and other markers of reactive astrocytes, previously characterized in MS (Girgrah et al., 1991; Ponath et al., 2018) (Fig. 3K,L). It is also marked by expression of tenascin C (*TNC*) (Fig. 3L), which encodes an extracellular matrix protein that was reported to support astrocyte morphological changes during development and activation (Roll and Faissner, 2019; Wiese et al., 2012). Interestingly, we also detected a strong astrocytic upregulation of *SHISA9* gene (Fig. 3M,N) that codes for CKAMP44 AMPA receptor auxiliary subunit (Farrow et al., 2015; Khodosevich et al., 2014), and has not been previously reported being expressed by human astrocytes. Neuronal death in MS leads to an abundance of cell-free glutamate, and its excitotoxicity has been shown, in turn, to be a major cause of neurodegeneration in MS (Pitt et al., 2000). AMPA receptors are the major glutamate receptors in the brain, and upregulation of a key AMPA receptor modulating protein in astrocytes, CKAMP44, might be associated with astrocytic transition to disease-state, and may have further implications for glutamate regulation in MS (Fig. 3M,N). Overall, these findings suggest enhanced activation of astrocytes in MS, accompanied by modification of the extracellular matrix and increased responsiveness to glutamate excitotoxicity - a hallmark of MS (Pitt et al., 2000), all likely contributing to inflammation.

Cluster-free analysis can reveal changes in subpopulations below the resolution of the utilized annotation. The investigator can then either adjust the resolution of the annotation to explicitly separate such subpopulations, or interpret the changes based on local gene markers or other properties as we have done above. The resolution and accuracy of cluster-free analysis is generally limited by the quality of the dataset alignment, and in case of embedding-based estimates, by the quality of the utilized embedding (Chari et al., 2021).

### Cluster-based expression shifts

Complementary to the compositional shifts are the changes in the expression state of different cell types. Even if the composition of the tissue remains largely unperturbed, the disease or condition being investigated can alter the expression state of the cells. The cell types that exhibit the greatest expression shifts between conditions are of interest as they are also likely to have altered their phenotype. While conceptually, performing such expression shift prioritization is simply a matter of comparing cross-condition expression distances for each cell type, the sparse and high-dimensional nature of the scRNA-seq data poses some challenges. To our knowledge, only one published method, Augur (Skinnider et al., 2021), aims to perform such analysis. Augur uses a random forest classifier to measure separability of two conditions, providing an intuitive metric as well as ranking of genes contributing to this separability. On the downside, Augur does not take into account variability between samples, which, as we will show later, can be crucial for robustness. We developed a different approach, focused on the analysis of pairwise distances between samples within each cell type.

We start by estimating correlation distances between all pairs of samples using pseudo-bulk expression profiles. We find that such distances are greatly affected by cell-type specific covariates, such as the average cell depth or the number of cells, presenting a key challenge for comparisons across cell types (Fig. S1A,B, Supp. Note 1). To overcome these biases, we normalize the distances between conditions by subtracting the average of distances within each condition (Fig. 4A-C). Furthermore, the normalization centers the distances by a background distance distribution obtained by randomizing condition group assignment across samples (see Methods). This procedure largely attenuates biasing effects of technical factors and provides a uniform way to assess statistical significance of the distances observed between case and control conditions (Fig. S1A-C). Applying the developed method to the PF dataset revealed that transitional AT2 and AT2 cells have one of the largest expression shifts (Fig. 4D). This might indicate that transitional AT2 cells (i.e. those that differentiate towards AT2 and AT1 cells) are activated to replenish AT2 and AT1 that die upon lung injury and are dramatically reduced in PF (Fig. 2H) (Abdelwahab et al., 2019; Lee et al., 2018). Among the datasets we have examined, the strongest shifts were observed in the squamous cell carcinoma (SCC) study (Ji et al., 2020), where the fibroblast population showed a prominent expression shift between the tumor and the adjacent normal samples (Fig. S1C). This difference likely captures the transition between healthy and cancer-associated fibroblasts (CAFs), which play an important role in formation of the fibrovascular niche in SCC (Ji et al., 2020). However, these differences are likely artificially increased by the procedures in the original preprocessing of the dataset (see Methods).

**Figure 4.**
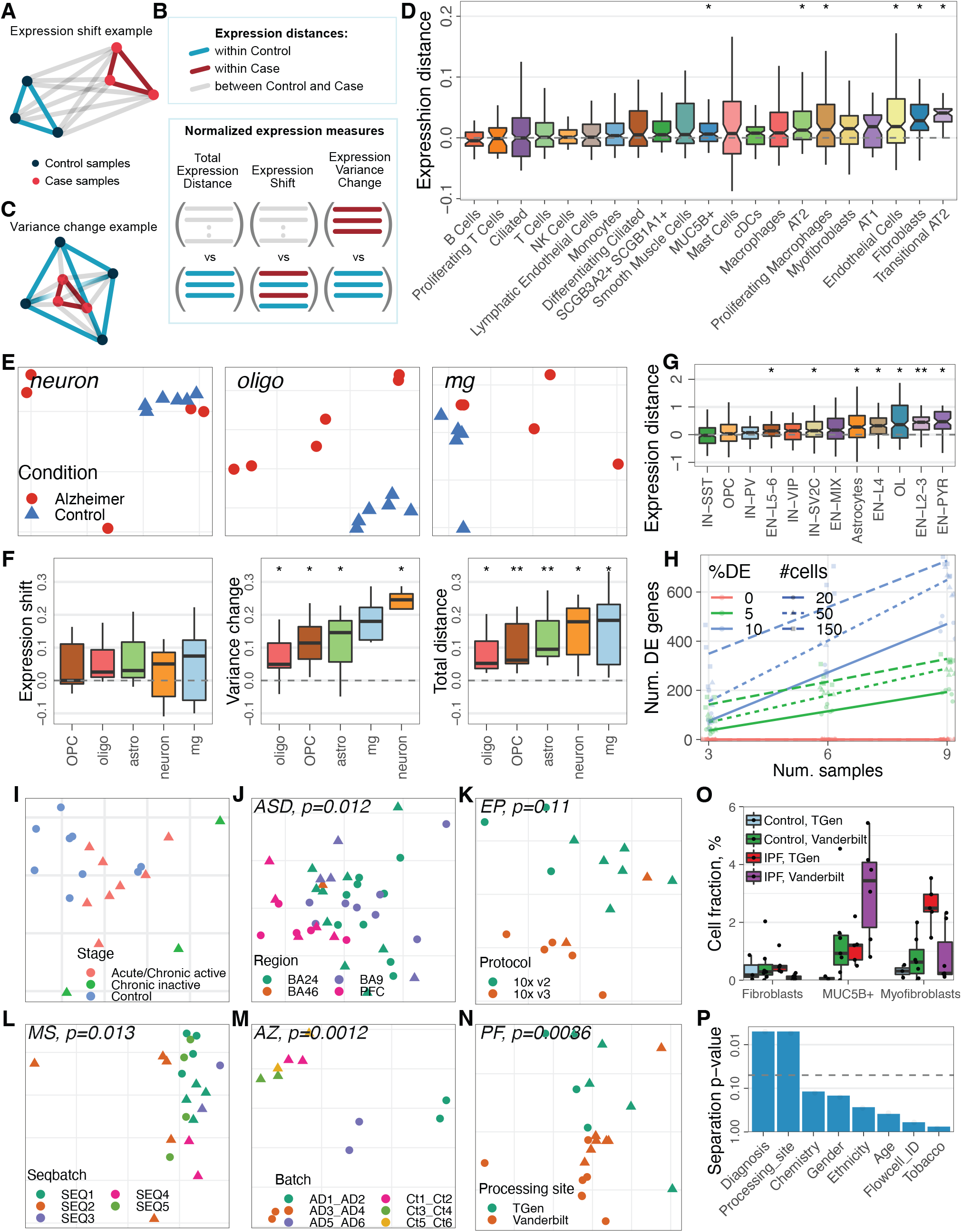
Cluster-based analysis of expression change magnitudes. (A-C) Schematic expression shift magnitude estimation. (A) For each cell type, the expression distances are estimated within (blue, red) and between (grey) conditions. (B) These distances can be normalized in different ways to capture different types of scenarios. Within each cell type, expression is summed by samples, and all pairwise distances between samples are estimated (A). The distances are normalized to adjust for potential confounders, with different normalizations capturing different types of magnitude (B). Normalizing pairwise distances between conditions by the median of within-condition distances captures “expression shift” between conditions (middle). Normalizing within-Case distances by within-Control distances, can capture change in the variability of expression (right, also see panel C). Finally, while normalizing between-condition distances between by within-Control distances captures both types of changes (“total expression shift” - left). (D) Expression shifts between Control and IPF conditions (y-axis) are shown for different cell types in the PF dataset. Boxplots represent distribution of normalized pairwise distances (B), where each observation represents an expression shift for a pair of samples from different conditions. Significance levels, estimated by a permutation test (with BH adjustment), are indicated by star symbols on the top. (E) Pairwise expression distances between samples are shown for three cell types from the Alzheimer dataset, using MDS embeddings. Each dot is a sample, with colors and point shapes corresponding to the sample condition. The ‘neuron’ cell type is an example of a variance change between conditions. (F) Different types of expression changes (see B) are shown for the Alzheimer dataset. (G) Expression shifts in the MS dataset (same format as D), estimated using 500 most differentially expressed genes. (H) The number of DE genes is a popular metric for measuring the extent of expression changes within cell type. The plot shows the dependency of this metric (estimated using DESeq2 with BH adjustment, y-axis) on the number of samples (x-axis), and number of cells (line type) for a simulated dataset. Different colors represent different fractions of simulated DE genes. Each dot corresponds to an independent simulation, repeated 5 times for each set of parameters. Both covariates strongly affect the metric and, hence, must be controlled for. (I) Visualization of sample structure using MDS embedding (x- and y-axes) of pairwise expression distances for the EN-L2-3 cell type in the MS dataset. Samples (dots) separate based on the disease stage (color). (J-N) Visualization of covariate structure in the expression space for different datasets, using MDS embeddings. Each point is a sample with shape representing condition. Color is used to visualize different covariates: Capture Batch for MS (J), Region for Autism (K), Batch for Alzheimer (L), Protocol for Epilepsy (M) and Processing site for PF (N). P-value for statistical separation of each covariate is shown on the top (see Methods). (O) Abundance of different fibroblast populations in the PF dataset is shown across conditions (x-axis) and processing sites (color). Each dot represents a sample. There is a clear difference in the direction of change depending on the processing site. (P) Separation p-values (y-axis, log-scale) for different covariates (x-axis) in the PF dataset. The horizontal line corresponds to the significance level of 0.05. P-values were adjusted using BH correction. Significance codes: \**p <* 0.05, ** *<* 0.01, * * * *<* 0.001.

The expression shifts between conditions are often driven by a limited fraction of affected genes, and such signal may get diluted by high dimensionality of the expression space. Furthermore, most distance metrics, such as Euclidean and even Correlation distance, lose sensitivity in high-dimensional spaces (Kharchenko, 2021). Conventional scRNA-seq analyses typically reduce dimensions by selecting most variable (overdispersed) genes, and further focus on the most pronounced axes of transcriptional variation using PCA or other dimensionality reduction techniques (Kharchenko, 2021). Such an approach, however, has limited applicability in the comparative setting. Running either overdispersion prioritization or dimensionality reduction on the entire dataset will bias the resulting space towards genes distinguishing individual cell types and subpopulations (i.e. marker genes), making the resulting space highly biased for assessment of expression shifts. Even if applied within the individual cell types, these steps would improve sensitivity only in situations when the transcriptional difference between the conditions being compared is the main driver of variation, which is often not the case, as we will illustrate in the next section.

To perform dimensionality reduction in a manner that would not bias dimensions towards specific cell types, we select genes that are most differentially expressed between conditions. Gene selection and the subsequent PCA dimensionality reduction is performed separately for each cell type. Because this procedure is designed to focus on the difference between conditions, the magnitude of the distances in the resulting space has to be interpreted relative to an expected background distribution, generated empirically by repeating the same procedures while randomizing sample assignment to groups (see Methods and Supp. Note 1). This “focusing” procedure greatly improves sensitivity: in the MS dataset, no significant shifts were detected without it, but when using 500 top-DE genes, significant shifts were detected for seven cell types (Fig. 4G). This focused analysis revealed significant shifts in gene expression for the excitatory neurons in L2-4, which were also shown to be reduced in abundance in MS by the compositional analysis (Fig. 2J). Such changes suggest vulnerability of upper layer cortical excitatory neurons to neurodegeneration. Curiously, we also identified a significant change in expression shift in astrocytes. While the original study did not report large expression changes in astrocytes, the impairment of astrocytes in MS was noted in the same study by smFISH, and the involvement of reactive astrocytes in tissue damage was further investigated. This example illustrates how expression shift analysis can effectively prioritize cell types that are likely to be functionally relevant for disease pathogenesis and could be further studied by histology and functional analysis.

In a simplest case, the impact of a disease on a cell type may result in a shift of expression state along some characteristic direction across all of the samples (“mean shift”, Fig. 4A). The impact of the disease in different samples, however, may be heterogeneous, and result in shifts of disease samples along different directions, effectively increasing variability of the expression states in the disease condition (“variance change”, Fig. 4C,F) (Batiuk et al., 2020). Such situation is likely to be characteristic of complex spectrum disorders, such as schizophrenia and autism, where understanding of multi-directionality of transcription shifts could help to distinguish subgroups of patients. Various scenarios can be quantified by contrasting the distances within and between conditions in different ways (Fig. 4B). We find that normalization of between-condition distances by the mean of distances within both disease and control conditions effectively captures the “mean shift” behavior (see Supplementary Note). On the other hand, normalizing between-condition distances solely by distances among the control samples captures a change in the “total expression distance”, which may reflect either presence of mean shift or increased variance in disease (Fig. 4B). Finally, the extent of “variance change”, can be evaluated by contrasting the distances within disease condition to those within control condition. Adjustment and statistical evaluation of all three measures follows the same logic, comparing the observed distances with the cell-type specific empirically estimated background distributions generated by randomizing sample assignments to conditions. For example, analyzing data on Alzheimer’s disease patients, we find that neurons as a whole do not show a significant mean shift, but instead show significant “variance change”, and as a result increased “total expression distance” (Fig. 4E,F). Most of the other cell types, such as astrocytes show both.

A popular metric for ranking the most impacted subpopulations relies on the number of differentially expressed (DE) genes (Grubman et al., 2019; Schirmer et al., 2019). The main caveat of such a metric is its dependency on the statistical power of DE tests, which varies depending on the number of cells and samples (Fig. 4H, Supp. Note 1). These critical covariates also vary between cell types, which can lead to notable biases. In general, more abundant cell types will typically yield a higher number of DE genes. While the dependency on the number of cells is often controlled by subsampling (Grubman et al., 2019; Schirmer et al., 2019), the variability of the number of samples in which the cell type is detected in sufficient numbers is often ignored (Fig. S1E). Controlling for the number of samples is also much harder in the studies with limited numbers of samples, as setting a minimum number of required samples too low would drastically increase variability of the metric, while setting the threshold too high would exclude many cell types that are not detected in a sufficient number of samples (Fig. S1F). Applying subsampling for both cells and samples, we found that on the SCC and MS datasets the results were relatively stable and consistent with the expression shift analysis (Fig. S1G,H). The main observed difference in the MS dataset was in the EN-PYR cell type, which was ranked first by the shift magnitude, but was omitted from the DE analysis because it was detected in less than six samples per condition (at minimal 30 cells per sample). Analysis of the PF dataset, in contrast, showed that setting the number of samples above 3 omitted >70% of cell types, while setting it to 3 or less resulted in too much uncertainty to distinguish any cell types as having significantly more DEs (Fig. S1F).

### Analysis of sample heterogeneity

While case-control studies focus on the condition-associated differences, there are usually other factors that affect the compositional and transcriptional features of the samples. These may include technical batches, patient’s disease stage, age, etc. For a given study design, typically many such sample covariates are tracked. Monitoring how these factors influence transcriptional and compositional characteristics of the samples is crucial for interpretation and reproducibility of the results. A simple way to explore the relationship of covariates with the expression state for a particular cell type is to visualize covariates on a plot of inter-sample expression distances. We used multidimensional scaling (MDS) for such visualizations. To assess whether a given covariate separates sample collection in a statistically significant way, we represented sample distances as a fully-connected graph and evaluated variation of the covariate on that graph (Shuman et al., 2013). The statistical significance of such a graph variation statistic was then assessed based on random permutations of covariate values across samples (see Methods, Fig. 4P). Visualizing disease stage in MS data for the L2-L3 EN subtype shows clear separation of acute and chronic stages (Fig. 4I), confirming the findings of the original publication (Schirmer et al., 2019). Furthermore, the expression distances can be aggregated across all cell types to visualize effects manifested in the full dataset in the same fashion. Technical batches and other experimental design factors often have substantial impact on the data, and if left unchecked can confound the interpretation of the results. Examination of the six datasets that reported sample metadata, revealed that technical covariates were responsible for a major portion of inter-sample variation in five of them. These covariates included sequencing batch – Alzheimer’s disease (AD) (Grubman et al., 2019) and MS (Schirmer et al., 2019), brain region – Autism (Velmeshev et al., 2019), 10x Chromium chemistry – Epilepsy (EP) (Pfisterer et al., 2020), and processing site – PF (Habermann et al., 2020) (Fig. 4J-N). Presence of such a strong separation indicates the need to control for these covariates, for instance by stratifying the data by the corresponding design factor, and ensuring that the experimental design is not confounded (e.g. each level of covariate is sufficiently well represented in both conditions being compared). Such stratification was done in the original PF, EP and Autism publications, but not in MS and AD.

Similar strategy can be followed to explore potential association of covariates with compositional variation between samples, simply by using ilr reduced space distances as a basis for MDS plots (Fig. 2B). In the PF dataset, for example, such analysis reveals striking differences in cell type composition between the processing sites (Fig. 4O and S2B), reflecting site-specific sample handling (Habermann et al., 2020). Variations in tissue processing protocols can distort cell type composition (Denisenko et al., 2020; Slyper et al., 2020), but in case of the PF study, CD45+ and CD45-cells were explicitly separated and re-mixed at pre-defined proportions at one of the processing sites. The analysis of cell abundance on the full PF dataset, therefore, is likely biased. Limiting the analysis only to the one site that did not perform such rebalancing reduces the number of samples and the resulting statistical power (Fig. S2A). In contrast to the full PF analysis, however, the analysis of a single site shows all subtypes of fibroblasts being overrepresented in PF, with Myofibroblasts being the most affected, which is consistent with existing knowledge of PF impact (Desai et al., 2018; Morse et al., 2019).

### Differential expression analysis

Functional interpretation of the expression changes in specific subpopulations ultimately relies on the analysis of differentially expressed genes. While many single-cell differential expression (DE) methods have been designed for comparison between subpopulations (Soneson and Robinson, 2018), here the aim is to identify genes differentially expressed within the same cell type between conditions (e.g. disease vs. control). The most common approach for carrying out such tests relies on “pseudobulk” expression profiles, which for a specified cell subpopulation aggregate molecules for a given gene across all cells in each sample (Crowell et al., 2020). This transforms the problem into a familiar setting in which multiple bulk RNA-seq profiles need to be compared between conditions. Such tests can be carried out using well-established standard packages, such as DESeq2 (Love et al., 2014), edgeR (Robinson et al., 2010) or limma-voom (Law et al., 2014). These common tools also provide standard means to control for the desired experimental covariates, such as age, or post-mortem interval. The pseudobulk approach, in essence, uses scRNA-seq to perform *in silico* purification of a specific subpopulation, discarding expression variability of the resulting cells within each sample. This is a reasonable approximation since for a sufficiently granular cell annotation, the variability of cells within a sample from a given individual will be notably smaller than between individuals. More complex analysis methods such as MAST (Finak et al., 2015) and *muscat* (Crowell et al., 2020), have been proposed to assess both inter- and intra-sample variability for a chosen cell subpopulation (Soneson and Robinson, 2018). Such methods, however, do not appear to show notable advantages in comparisons of averages (Crowell et al., 2020; Soneson and Robinson, 2018), and in practice are considerably slower than the pseudobulk alternative (Soneson and Robinson, 2018). It was also shown that the existing single-cell methods are usually biased towards highly-expressed genes (Squair et al., 2021), and often fail to accurately control the false-discovery rate (Zimmerman et al., 2021). In the context of multi-sample single-cell analysis, such problems may be amplified.

Since the aim of the differential test in a comparative setting is to identify genes that distinguish samples from the two conditions, we evaluated the sensitivity of the reported DE gene lists with respect to the exact set of patient samples included in the panel. We tested two approaches to threshold for DE genes: by taking genes passing a particular P value cutoff, or by taking top N genes. We then performed the leave-one-out (LOO) procedure on the samples (i.e. iteratively omitting one of the samples), and quantified the stability of the resulting DE gene sets using Jaccard coefficient (Fig. 5A,B). For all of the DE methods examined, selecting DE genes by top N resulted in a more stable gene set, however even the top N results showed only moderate stability, with Jaccard coefficient around 0.6 for top 100 genes (see Fig. S3A for other values of N). The stability of the DE results showed systematic variation across cell types (Fig. 5D) and across datasets (Fig. 5E). The stability was also low when summarizing DE genes in terms of enriched GO pathways, though GSEA performed better than hypergeometric enrichment tests (Fig. 5C).

**Figure 5.**
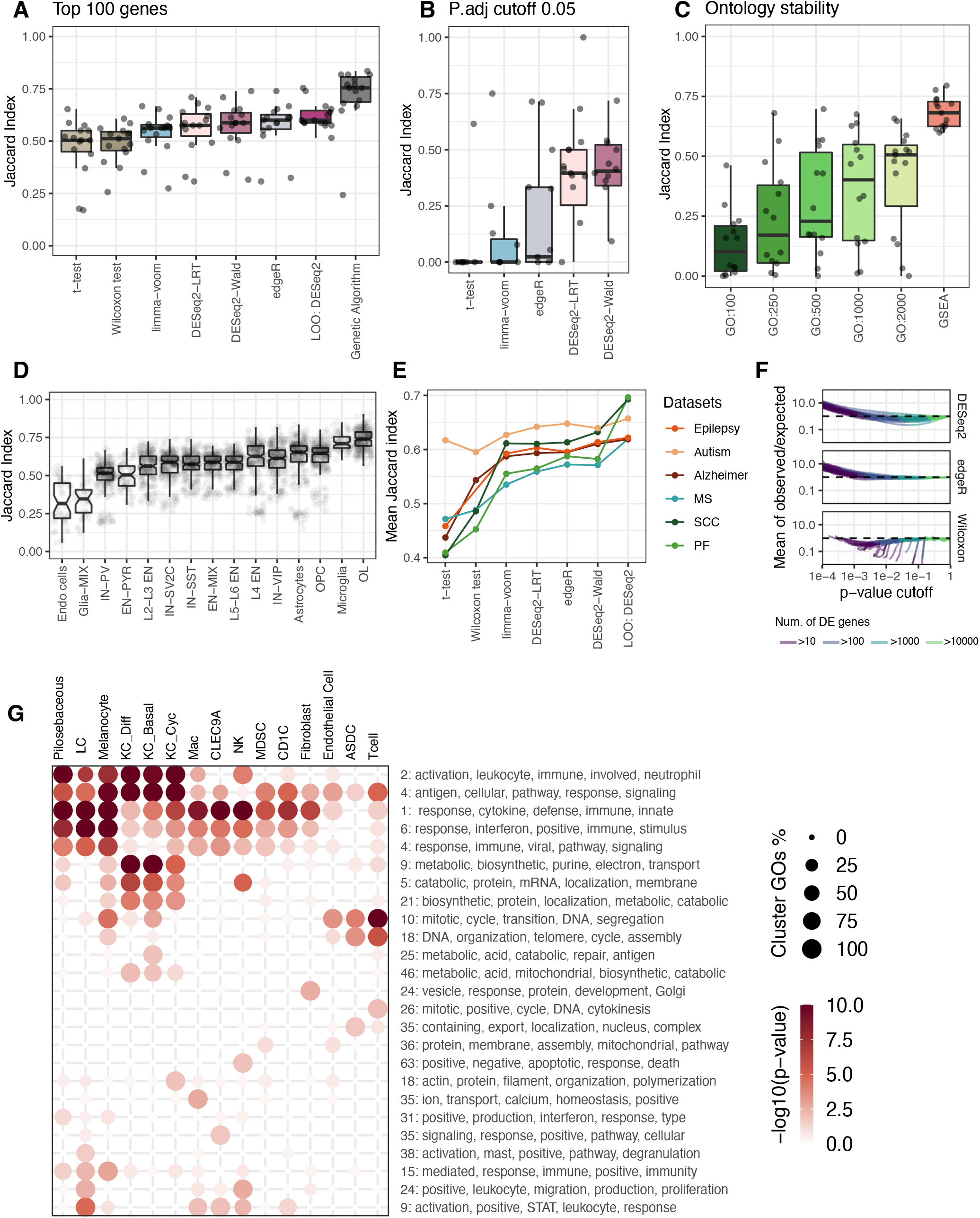
Differential Expression and Gene Ontology analysis. (A) The stability of differential gene expression results across leave-one-out iterations is assessed using Jaccard Index (y-axis) between “top 100” DE gene sets estimated for different methods (x-axis) on the MS dataset. Each dot corresponds to a specific leave-one-out configuration. (B) Analogous to (A), the plot shows stability of the DE gene sets, selected based on the p-value threshold (P < 0.05, adjusted for multiple hypothesis). (C) Jaccard Index between the sets of enriched Gene Ontology (GO) terms (y-axis) is shown for different estimation methods (x-axis). The methods include running GO on different number of top DE genes (100-2000) and for GSEA. (D) Stability of the “top-100” DE gene set is shown for different cell types of the MS dataset. (E) Average stability across cell types is shown for different datasets (color) and different test methods (x-axis). (F) The number of DE genes reported by a test (DESeq2, edgeR, Wilcoxon) for different levels of statistical stringency (p-value cutoff, x-axis) is shown relative to the number expected from a uniform p-value distribution (observed/expected ratio, y-axis). The DE tests were carried out on the PF dataset, randomizing sample assignment between conditions. Each curve shows the average observed/expected ratios for one cell type. Both DESeq2 and edgeR report over-abundance of DE genes at high confidence levels. (G) Dotplot visualization of Gene Set Enrichment Analysis of the SCC dataset. Each column corresponds to a cell type and each row to a cluster of GO terms, estimated by first grouping terms with similar sets of enriched genes, and then by the patterns of their expression across cell types. Dot color shows mean significance level calculated on log scale across the group for the given cell type. Dot size indicates the fraction of terms from the cluster which are enriched in the corresponding cell type. The row labels show the number of GO terms in each cluster, followed by five most frequent words in the contained GO terms.

As poor stability of the DE results poses a problem for downstream interpretation, we looked for potential explanations and approaches to mitigate this instability. Simple adjustments, such as exclusion of lowly-expressed genes (Squair et al., 2021) provided only minor improvements (data not shown). We have also explored whether the stability can be improved by reporting a “robust” subset of DE genes, that is those genes that are most frequently reported as DE genes when performing leave-one-out perturbations of the sample collection. We found, however, that such meta-approach provides only moderate improvement to the stability (Fig. 5A – “LOO”). Finally, we postulated that poor stability may be due to some, potentially unobserved, sample covariate. Such covariate could be, for example, a sample viability characteristic such as RNA integrity, or a design characteristic such as patient age. To test this, we used machine learning to fit a latent factor across samples that optimized stability of DE results for a given cell type under the sample leave-one-out procedure (Fig. 5A - “Genetic Algorithm”, see Methods). We found that such latent factors optimized for different cell types tended to be similar (Fig. S3B), and that factor optimizing stability for one cell type, on average, significantly increased DE stability for all the other cell types (Fig. S3C). The pathways enriched in the DE results remained similar when performing such a latent factor correction (Fig. S3D,E), suggesting that the latent factor was capturing a technical covariate. However, the factors did not strongly correlate with the covariates captured in the study design (Fig. S3B).

To understand the reasons behind moderate DE gene set stability, we re-evaluated the basic characteristics of the DE methods under a permutation test, that is examining whether the distribution of P values reported by a method follows uniform distribution under the null hypothesis. Specifically, we randomized samples across treatment across sample groups (i.e. randomized disease/control assignment) and calculated the ratio of observed to expected fraction of genes below different P value thresholds. Specialized RNA-seq DE tests showed a consistently biased trend which over-estimated the number of statistically significant DE genes (Fig. 5F). This behavior was more pronounced for some datasets than others (Fig. S3F,G). It should be noted that under such settings even the standard Wilcoxon rank sum test showed deviations from the null, though of lower magnitude and often in the opposite direction – underestimating the number of highly differential genes (Fig. 5F, S3F). Overall, our results illustrate that in the context of comparative analysis, the existing DE methods lack robustness and can overestimate the statistical significance of top DE genes. As a result, the effective stability of such results with respect to the exact set of samples being analyzed will be moderate. The limitation likely stems from under-estimation of the inter-sample variability and can be alleviated by larger panel sizes and improved statistical models.

The functional interpretation of the DE results commonly relies on analysis of pathway enrichment. Cacoa package implements several features to facilitate this, including interactive tables that allow for sorting, filtering and enrichment testing of DE genes reported for different subpopulations. The package also implements functions for providing overviews of pathway enrichment results using hierarchies or pathway clusters. As GO results can include many closely-related terms, the overview plots can collapse GO terms based on gene and pattern similarity. For the SCC dataset, such overview shows a number of relevant broad and cell type-specific changes, including i) modulation of migration of immune cells, such as NK cells, macrophages, and Langerhans cells, ii) enhanced proliferation/cell cycle entry of T cells, iii) increased metabolism in keratinocytes (KC_Diff, KC_Basal), and finally, iv) a large cluster of immune response pathways encompassing a number of myeloid cell types, such Langerhans cells, macrophages, and dendritic cells (*CLEC9A* and *CD1C*), as well as NK cells, Pilosebaceous keratinocytes, and Melanocytes (Fig. 5G).

### Cluster-free expression shifts and differential expression programs

Quantification of expression shift magnitude can be carried out in a cluster-free manner. We approached this by analyzing the neighborhood of each cell and estimating pairwise distances to the cells from the other samples found in the neighborhood. The same p-value adjustment procedure as for the cluster-free composition shifts was applied (see Methods). Generally, cluster-free shifts closely match cluster-based results, though typically show lower statistical power (Simulation Note). As expected, however, they can also reveal finer details. For example, we find that within the Endothelial cells, the Pulmonary-venous (CPE+) and the Arterial (*DKK2*+) subtypes (Schupp et al., 2021) exhibit the highest magnitudes of transcriptional change in PF patients (Fig. 6A-C). Various studies have suggested that changes in endothelial cells can be due to endothelial-to-mesenchymal transition (Leach et al., 2013). So, we tested two markers of this transition (*FN1* and *S100A4*) (Leach et al., 2013), and found that PF patients showed significant upregulation of *FN1* in both Pulmonary-venous and Arterial cells, while *S100A4* was upregulated in Arterial, General capability cells and Aerocytes (Fig. 6D,E).

**Figure 6.**
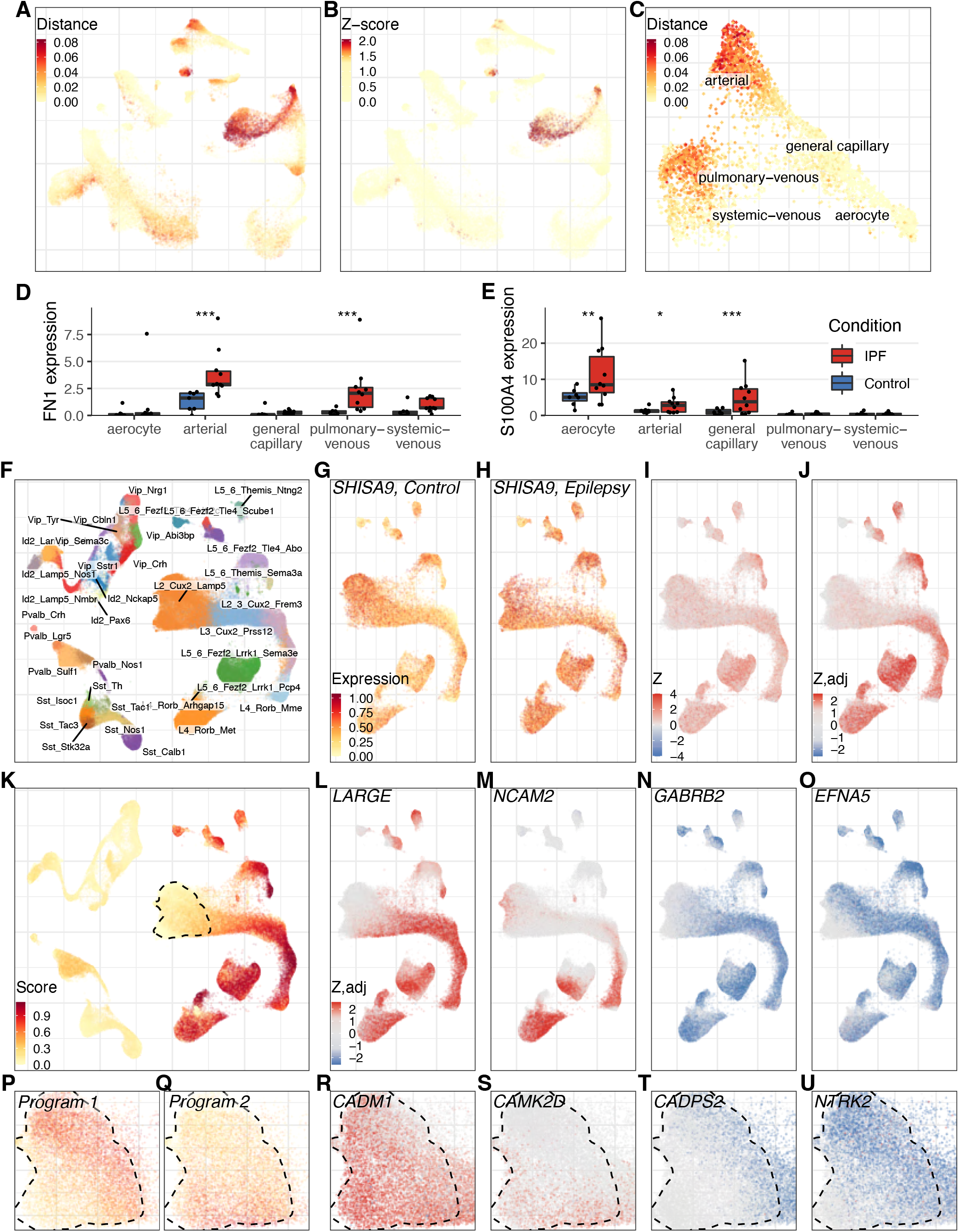
Cluster-free gene expression analysis. (A-C) Cluster-free analysis of expression shifts in the PF dataset. (A) shows expression shift magnitudes (color) on the whole dataset with (B) showing adjusted statistical significance levels. As the Endothelial cells show heterogeneous signal, we re-annotated them based on known markers (Fig. 3I) and carried out focused re-analysis of the Endothelial cell subpopulations alone (C). (D-E) Normalized expression of Endothelial-to-Mesenchymal transition markers (Leach et al., 2013) with the asterisks showing the significance levels (DESeq2 test with BH adjustment: \**p <* 0.05, ** *<* 0.01, * * * *<* 0.001). (F) Annotated UMAP for the Epilepsy dataset. (G-J) Visualization of cluster-free differential expression of SHISA9 gene in the Excitatory neurons of the Epilepsy dataset. Embedding is colored by the SHISA9 expression level in Control (G) and Epilepsy (H), difference statistics (I), and statistical significance of the expression change (J). (K) Visualization of the global score (color) for the gene program that involves most of the Excitatory neurons. The contour outlines the L2_Cux2_Lamp5 cell type. (L-O) Significance of change of four genes (top-left corner) that contribute to the program shown in (K), shown using adjusted z-scores (color). (P-Q) Local gene program scores for two most pronounced programs, estimated within the L2_Cux2_Lamp5 cell type of the Epilepsy dataset. (R-U) Adjusted z-scores are shown (color) of four genes (top-left corner) that contribute to the programs shown in (P) (R-S) and (Q) (T-U).

We also extended DE between conditions so it can be carried out in a cluster-free manner. To do so, we quantify the expression difference between conditions in the neighborhood of each cell as defined by the joint dataset alignment graph. For each gene, we quantified the expression difference as a Z score statistic (Fig. 6G-I, see Methods), and then evaluated statistical significance using the same empirical randomization procedure as described for the cluster-free compositional analysis (Fig. 6J). These Z score estimates can be used to detect *gene programs* – a combination of similarly affected DE genes and the groups of cells where their difference is most pronounced. Detection of such gene programs is analogous to the traditional bi-clustering problem (Padilha and Campello, 2017; Pontes et al., 2015), however existing algorithms that we tested (Clevert et al., 2015; Duarte and Mayrink, 2019; Hochreiter et al., 2010; Li et al., 2009; Secchi et al., 2016), scaled poorly with the number of cells in the datasets (data not shown). Instead, we used traditional Leiden or PAM clustering of genes, followed by a search of cell subpopulations where the mean Z score of the cluster was high (see Methods).

This gene program approach is able to find DE patterns shared by multiple subpopulations of cells (Fig. S4A). For example, in the Epilepsy dataset, we find a program that involves many subtypes of principal neurons in epileptic cortex (Fig. 6F,K). This program is driven by genes enriched in cellular development GO terms (Fig. S4B). It also captures upregulation of neuronal cell adhesion and neurotransmission genes (Fig. 6L-O), including *SHISA9* (Fig. 6G-J) - a gene encoding critical AMPA receptor modulator that was previously shown to be upregulated in epilepsy across layers of principal neurons (Pfisterer et al., 2020), and was noted above for upregulation of its expression in an expanding astrocyte population in MS (Fig. 3M,N).

Gene programs can be also used to examine expression differences within specific cell types of interest, potentially going beyond the resolution of annotation. For example, focused re-analysis of the L2_Cux2_Lamp5 Excitatory neurons showed separation of two DE programs within this population, which also matched subpopulations detected by the cluster-free composition analysis (Fig. 6P,Q and S4D,E). The first program was associated with increased expression of cell adhesion and neuronal plasticity genes, such as *CADM1* and *CAMK2D* (Fig. 6R,S), with CAMK2D known to be involved in plasticity and neurite outgrowth of principal neurons (Zalcman et al., 2018). The second gene program was enriched for neurotransmission and developmental GO terms. It involved downregulation of neurotrophin signaling modulators (*NTRK2* and *CADPS2)* (Fig. 6T,U), which are coding for the major receptor for BDNF and NT-4/5 neurotrophic factors (NTRK2/TrkB protein) (Pezet and McMahon, 2006), and regulator of neurotrophin (CADPS2) (Sadakata et al., 2004).

### Iterative functional interpretation

The panel of analysis methods implemented in Cacoa enables interactive analysis of scRNA-seq data, facilitating an iterative process of explorative analysis, hypothesis formulation, and focused follow up. In the case of the PF dataset, for example, our initial Cacoa analysis confirmed the major findings reported in the original publication (Habermann et al., 2020), such as the notable expansion of the KRT5-/KRT17+ epithelial cells, as well as their modulation of the extracellular matrix organization in PF (Fig. 2H and 7A). The results extended the observations of the original publication, revealing significant disease-associated changes in the subtype composition and gene expression in the Endothelial and Immune compartments (Fig. 2H, 3H, 4D, 5C), and providing further resolution in the complexity of changes in Epithelial cells (Fig. 2H, 4D).

**Figure 7.**
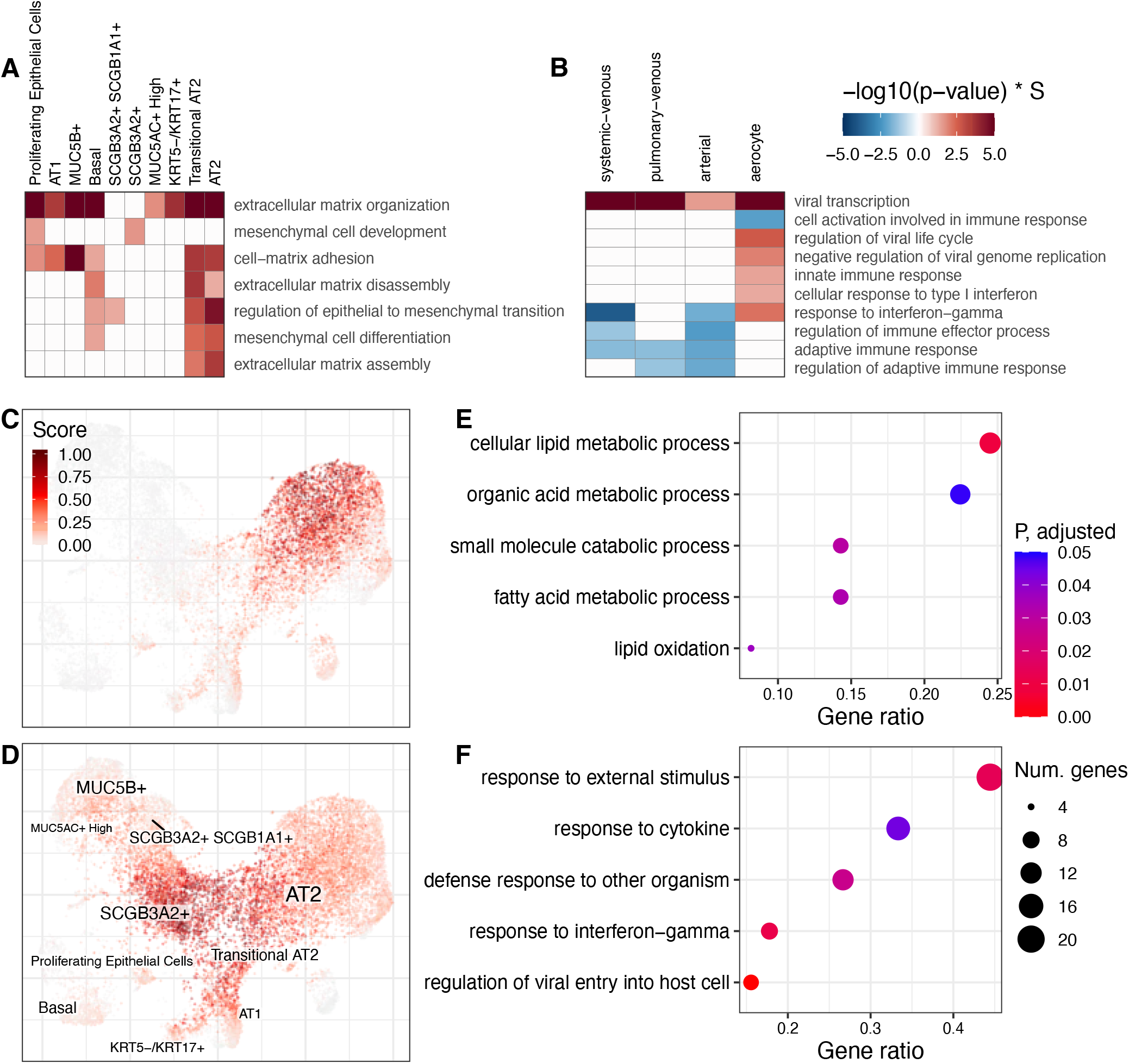
Functional interpretation. (A-B) GO heatmap for different groups of ontologies in the PF dataset. Each row corresponds to a cluster of ontologies identified based on similar genes (row name shows medoid of this cluster), each column is a cell type and colors show significance of the corresponding medoid ontology, found by GSEA. Only significant terms are shown (p-value < 0.05 after BH adjustment). Red colors correspond to up-regulated pathways, and blue colors show significance of down-regulated ones. (A) Extracellular matrix and mesenchymal transition ontologies across Epithelial cell types. (B) Virus and immune response ontologies across Endothelial cell types. (C-D) Scores of two cluster-free gene programs, shown on the focused UMAP embedding of the Epithelial cells in the PF dataset. The program shown in (C) involves a subset of AT2 cells. The program shown in (D) captures the transition point from AT2 and SCGB3A2+ types to AT1 and KRT5-/KRT17+. The color represents the program score in the given cell. See Figure S6C for the annotation of the UMAP embedding. (E-F) GO terms (y-axis), enriched in the programs shown in (C) and (D), respectively. The x axis shows the fraction of expressed genes from a GO term that were part of the program. Dot size indicates the number of such genes, and the color shows adjusted p-value (BH adjustment). To reduce redundancy, GO terms were clustered based on the composition of the genes enriched in them, and for each cluster, a GO with the most representative name was shown (see Methods).

To illustrate follow up analysis we focused on the AT cells, which together with the SCGB3A2+ and KRT5-/KRT17+ cell types exhibited the largest magnitude of expression changes in PF. Cluster-based DE analysis showed that AT2 cells have a strong upregulation of extracellular matrix organization pathways (Fig. 7A), indicating that together with previously reported KRT5-/KRT17+ cells (Habermann et al., 2020), AT2 cells are likely involved in the remodeling of the extracellular matrix in PF. We also looked for potential explanations to a marked reduction in the number of all annotated AT subtypes (Fig. 2H), and found that AT1, AT2, and Transitional AT2 cells upregulated cell death pathways. In particular, the GO term “epithelial cell apoptosis process” was upregulated in all AT cells, but not other cell types (Fig. S5A), reflecting specific prevalence of apoptosis of the AT compartment, consistent with previously published observations (Abdelwahab et al., 2019; Lee et al., 2018). Additionally, we noted a complex inflammatory response in epithelial cells, leading to an increase in the expression of inflammatory pathways in AT2 and SCGB3A2+ cell types, while MUC5A/B+ and KRT5-/KRT17+ cell types showed decreased expression of inflammatory pathway genes (Fig. S5B). Cluster-free DE analysis of AT cells additionally revealed two gene programs, capturing PF-associated transitions within epithelial cells (Fig. 7C-D, S5C). The first program (Fig. 7C) likely represents transition from immature differentiating/de-differentiating AT2 cells and mature homeostatic AT2 cells and enriched in genes involved in lipid metabolism (Fig. 7E), a key intracellular signaling in mature homeostatic AT2 cells (Gokey et al., 2021). The second program (Fig. 7D) corresponds to a central transitional state between all 4 major subtypes of epithelial cells - AT1, AT2, SCGB3A2, KRT5-/KRT17+, which is strongly associated with immune response (Fig. 7F). Thus, implementation of Cacoa allowed to leverage scRNA-seq data in PF and to reveal potential pathogenic mechanisms that occur in highly complex epithelial tissue in the lungs.

Previously cluster-free expression shifts showed that Endothelial cells have heterogeneous response with arterial and pulmonary-venous cell types being the most affected (Fig. 6C). In such situations it is useful to re-annotate the region of interest in higher resolution and run the cluster-based analyses on that. We have, therefore, split Endothelial cells into five subtypes (Fig. 6C) and reran the DE analysis on these subtypes. It showed that arterial (also highlighted before) and systemic-venous cells have the highest number of immune response related pathways, while aerocytes show several pathways related to viral life cycle (Fig. 7B). Overall, our data shows great applicability of Cacoa to identify molecular and cellular mechanisms that could underlie disease pathogenesis in complex tissues.

## Discussion

Single-cell studies across conditions have potential to provide insights into disease etiologies, reveal molecular mechanisms of pathogenesis, and suggest targets for improved diagnosis and treatments. Indeed, a number of such studies comparing healthy and disease tissue states have been published in the last few years, and their numbers are likely to increase in the coming years. It is, therefore, important to develop robust tools for contrasting conditions with scRNA-seq data. Only a handful of relevant tools have been brought forward recently, including MELD (Burkhardt et al., 2021) and Milo (Dann et al., 2021), which perform cell abundance analysis, Augur - a method to prioritize those cell types that are most responsive to biological perturbations in single-cell data (Skinnider et al., 2021), and scCODA - a Bayesian model to identify the effect of disease on cell type composition (Büttner et al., 2021). Cacoa brings together a broader collection of tests and diagnostic tools for comparative analysis, including both cluster-based and cluster-free methods, as well as compositional and expression shift perspectives.

In implementing such methods in Cacoa, we noted that many seemingly simple comparisons can exhibit systematic bias or instability due to technical factors, requiring more elaborate measures and statistical controls. For example, measures capturing the extent of transcriptional difference between conditions were heavily biased by the number of measured cells, coverage, and other technical characteristics (Supplementary Note 1). In this and other cases, we resolved the issues by generating cell-type specific background distributions through randomization of sample assignment across conditions. Such empirical distributions enabled both bias adjustment and evaluation of statistical significance. As we illustrated with the re-analysis of six published scRNA-seq datasets, these types of statistical considerations, implemented in Cacoa, can reveal previously unappreciated aspects of disease impact.

We were surprised to find particularly poor stability of differential expression analysis, which relies on well-established statistical methods for bulk RNA-seq comparisons. The principal aim of the comparative analysis is to identify differences between conditions that exceed inter-individual variation, and it appears that the existing DE methods underestimate variation when given pseudo-bulk profiles from different individuals. It is unclear whether this is due to the technical sources of variability, such as variation in the number of cells or coverage, or biological variation between individuals. The stability can likely be improved by increasing the number of samples in a comparison, but in currently realistic study designs, which typically involve ∼10 samples per condition, the existing DE methods do not provide reliable lists of DE genes. As we noted, simple filtering and ranking solutions have yielded only minor improvements in stability. A more adventurous approach of finding a latent factor across samples that optimizes DE stability did improve stability (Fig. 5A), however whether such “stabilizing” factors are biologically meaningful is difficult to assess. Notably, the stabilizing factors determined for different cell types appeared to be relatively similar (Fig. S3B), suggesting that instability is driven by sample-wide effects, and hence may be possible to correct effectively. The number of significant DE genes has also been widely used as a way of quantifying the extent to which different cell populations have been impacted by the disease. Such a measure, which depends on statistical power, is heavily biased by the number of cells observed for different cell types, and hence some published studies have controlled for that. We show, however, that the number of DE genes is also biased by the number of samples in which the cell type appears – a factor that has so far been ignored. Consequently, we recommend using expression shift measures as a more robust readout for the prioritization of cell type impact.

The variability between samples can be often traced back to known technical or biological factors, such as processing batch, age, etc. Using simple MDS plots and tests, we indeed found that in many cases batch or other technical factors drove expression or compositional differences of the datasets more than the conditions the study was designed to compare. Especially when unbalanced, such factors can significantly distort the results, highlighting the importance of tracking and testing for association with technical and design factors, as was done for instance in (Batiuk et al., 2020). Further methodological developments will be needed to incorporate ability to control and correct for extraneous factors within the comparison tests themselves. Such corrections may also increase the sensitivity of the tests. For instance, removing genes associated with sex and batch from the MS dataset, improved the sensitivity of the expression shift analysis, detecting the same subtypes as were shown when focusing on the top-500 most changed genes, indicating improved sensitivity (Fig. S2C-E).

It is typically easier to compare conditions using cell type or cluster annotations. Such cluster-based analysis also yields better statistical power, however, can be sensitive to the choice of the annotation and cluster boundaries. Cluster-free methods can provide guidance in that regard, by highlighting subpopulations most impacted by conditions, so that the granularity of the annotation can be adjusted to match the resolution of the data. In general, the choice of the annotation granularity connects the two facets of the comparison – compositional and expression state differences. For instance, coarser annotations can conceal compositional differences, translating them into expression shifts (Fig. 1C). Conceptually, separating compositional and expression shift effects is appealing, however in practice the two are very difficult to delineate. Even in an idealized case of an expression shift, when the impact of a disease, uniformly alters the expression state of a specific subpopulation, the difference may be captured as a compositional difference if, for example, the dataset alignment and clustering step groups disease-impacted cells into a separate cluster. With our examples, we therefore tried to emphasize the need for iterative analysis, involving careful diagnostic of the observed differences, adjustment of the annotation resolution, and focused re-evaluation of differences within compartments of interest.

Overall, the Cacoa package provides a broad range of methods for contrasting conditions with scRNA-seq collections illustrated above, including expression and compositional difference perspectives, as well as both cluster-based and cluster-free approaches. Starting with Conos (Barkas et al., 2019) or Seurat (Hao et al., 2021) alignment objects, Cacoa provides user-friendly ways of generating plots and tables that can be used to interpret the biological changes between conditions. It also brings robust statistical tests and means to screen for potentially confounding factors. As scRNA-seq is rapidly becoming the method of choice for understanding the impact of disease and other perturbations on complex tissues, we hope Cacoa will facilitate such analysis and help to avoid problems stemming from statistical bias or instability.

## Acknowledgements

We thank Katarina Dragicevic there for providing extensive feedback on the package functionality, and Evan Biederstedt for assistance with package maintenance. P.V.K was supported by NIH 1U54HL145608. K.K. was supported by the Novo Nordisk Foundation Hallas-Møller Investigator (NNF16OC0019920) and Hallas-Møller Ascending Investigator (NNF21OC0067146) grants.

## Competing Interests

P.V.K. serves on the Scientific Advisory Board to Celsius Therapeutics Inc. and Biomage Inc.

## Methods

### Cluster-based compositional changes

We use compositional data analysis (CoDA) theory to catch the differences in cell-type compositions between two sets of samples (e.g., case and control). The core idea in CoDA is to transform non-independent proportions to the Euclidean space with independent coordinates. One of these transformations is called isometric log-ratio (ilr) transformation. We first describe the ilr transformation, and then how we employ it in the comparison of two groups.

Let consider the dataset of *m* samples, each represented with counts of *n* cell types, so that all counts form the matrix *x* of size (*m* × *n*). The matrix of counts can be transformed to the matrix of frequencies, *f*, dividing each row by its sum. We apply isometric log-ratio transformation (ilr) to each row of *f* based on a random sequential binary partition of *n* cell types (Pawlowsky-Glahn et al., 2015). In brief, we generate a random binary tree with *n* leaves randomly corresponding to cell types and assign a balance function to each inner node as proportional to the log-ratio of geometric means of cell type frequencies in the two subclades of the node. Mathematically this transformation of frequencies to balances is expressed as follows:

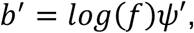

where *ψ* is the basis matrix of size (*n, n* − 1) encoding the structure of the binary tree, and 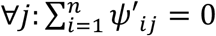. Instead of *n* cell type frequencies, (*n* − 1) balances are independent and can take any real values.

We perform PCA on matrix *b*′, obtain rotation matrix *R* and new balances *b* = *b*′*R*. Due to the ilr properties, new balances are also ilr-transformed frequencies, but with the basis *ψ* = *ψ*′*R*.

To catch the difference between two groups of samples, we applied linear discriminant analysis on balances. We estimate the following linear model: *g ∼ α* + *bw* + *ε*, where *g* is the group variable, *w* is the vector of linear influences of balances to groups to be optimized. After optimization, we divided estimates 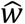 by standard deviation of corresponding balances to obtained loadings of balances to the group variable,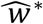. The corresponding loadings of cell types to the group variable we define as 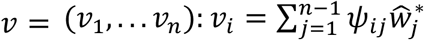.

After that we identified the loading level of reference cell types. While the reference cell type can be explicitly specified by the investigator, we also developed a way to identify the reference set automatically. Specifically, for each pair of cell types, we calculated the log-ratio value of their frequencies and tested its association with the grouping variable (e.g., case-control). If p-value of association was less than predefined cutoff (0.05), then changes in frequencies of these cell types were considered to be coherent. Following a greedy strategy, we grouped cell types with the coherent behavior until all remaining groups demonstrate pairwise significant differences (i.e. no coherent group pairs could be found). The group with the lowest mean value of loadings *v*_*min*_ was then assigned to eb the reference group. The loadings *v* were then normalized to the reference level with the subtraction of *v*_*min*_, so that mean value of loadings in the reference set was equal to 0.

To test for statistically significant changes in composition between case and control sets, we performed the analysis described above on the resampled datasets *t* = 1000 times. In each resampling, we randomly chose *m* samples with replacement, and then performed boobstrap resampling of cells in each sample. For a *k*-th resampling iteration, the procedure returns a vector of cell type loadings *v*^*k*^. To test for the difference in *i*-th cell type different between two groups of samples, we calculated empirical p-value as:

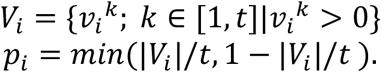

### Compositional space

Dimensional reduction of cell type compositional analysis was in ilr coordinates (balances). Specifically, PCA or CDA (canonical discriminant analysis) was performed on matrix *b*. In the case of two classes, CDA is equivalent the Fisher’s linear discriminant, and returns one direction in the space of balances that optimally separates the two classes. To visualize this separation, we used the CDA-produced direction and the first principal component estimated on the affine space orthogonal to the direction. The separation of more than two classes is visualized in two first CDA coordinates. Compositional distance, *D*^*coda*^, between samples is Euclidian distance in the ilr space.

### Compositional trees

A binary CoDA tree was constructed to visualize coherent compositional changes among different cell types. The hierarchical clustering was performed as described in the “Cluster-based compositional changes” section. Briefly, at each step, the algorithm jointed a pair of subclusters if the log-ratio balance between them demonstrates the lowest association with the sample group variable. Such greedy clustering approach groups most coherent cell types, separating most contrasting groups of cell types within the top splits of the tree.

### Cluster-free compositional changes

Estimation of cluster-free compositional changes goes in three steps. First, the densities of individual samples are estimated by using kernel density or graph density estimation (see below). Then, the difference of sample densities between conditions is estimated for each data point (by default, using Wilcoxon test statistics). Finally, condition labels per sample are permuted and the whole procedure is repeated 400 times to estimate significance of changes in each data point.

Densities of individual samples can be estimated using either cell graph or a low-dimensional (*e.g*., 2D) cell embedding. In the case of 2D embedding, this is done using kernel density estimates (KDE). For that, the whole 2D space is split into the grid of 400×400 bins, and the density of each sample is estimated for each bin using ks R package with subsequent normalization of values to unit sum per sample. The KDE bandwidth is estimated as 5% of the difference between 90% and 10% of the corresponding axis values. For estimation on the cell graphs, the algorithm operates on the level of individual cells instead of bins. It estimates graph densities following the method suggested in (Burkhardt et al., 2021). The algorithm converts sample labels to binary values using one-hot encoding and then applies graph heat filter to smooth this signal. As the result, it provides a vector of sample densities for each cell in the dataset.

When sample densities are estimated, the difference in each data point (bins for KDE and cells for graph density) is estimated as Wilcoxon test statistics. Other measures of difference can be applied, but Wilcoxon statistic showed the highest power on our tests. Wilcoxon test p-values could not be used directly, as it would require correction for multiple comparisons, and having so many correlated comparisons would result in extremely low power. So here and in other cluster-free methods we used the significance testing procedure proposed in Holmes et al. (Holmes et al., 1996) with several adjustments that are described below. For this procedure, we used cell graph for graph densities and 4-connected grid for KDE, applying the heat filter for signal smoothing and using winsorizing level of 1%. The number of permutations was set to 400 by default.

### Estimating significance for correlated observations

When doing statistical testing, most corrections for multiple comparisons assume independence of these comparisons. Thus, for the case of correlated observations they over-correct, which leads to reduction in statistical power. Moreover, when observations are highly correlated, such as cells with very similar expression profiles, they may be considered as replicate observations, and so having more of those should increase the power, which is the opposite of what the usual corrections do. One solution for that would be to split data into a small number of regions, which are considered independent from each other, while aggregating the signal within each region. A variation of this idea is implemented in the Milo package (Dann et al., 2021). An alternative permutation-based approach was suggested for image analysis in (Holmes et al., 1996). It can work in very broad settings, and below is the workflow adapted for our experimental setup.

The input to the procedure is a matrix of observations (*i.e*., cells in most cases), where each observation (row) is a vector of measurements in different samples (columns). Each sample has one of two condition labels, and for each observation we should be able to estimate a difference statistic between conditions. Example of such data would be sample densities per cell (or per bin) in cluster-free composition shifts. The difference statistic depends on the problem of interest, and the only requirement for it is to be approximately identically distributed in all data points. If it is not identically distributed, the test still controls for the false discovery rate (FDR) but has different power in different data regions. When the difference is estimated on the observations, the next step is to perform a large number of permutations of condition labels across sample, and to estimate the differences on the permuted labels. For each permutation, maximum of all differences is estimated, thus resulting in the vector of maximums across all permutations. Then, to control FDR at significance level of *α*, difference for each observation under real sample labels is compared to (1 − *α*) quantile of the permutation distribution, and differences above this quantile are considered significant.

We additionally adjusted this procedure to increase its sensitivity on scRNA-seq data. Given that the method is agnostic to the choice of the difference function, it allows for arbitrary post-processing transformations of the differences between conditions, as long as they are also performed consistently across all permutations. So, after estimating the difference statistics we first perform winsorization of the values by some small percentage (1-3%, depending on the problem). This step helps to stabilize the empirical FDR estaimtes, as the procedure operates on maximums and is very sensitive to outliers. Then, we perform smoothing of the difference statistics on the graph. Usually, it is performed on the cell graph, though the procedure can work with arbitrary graphs. The smoothing is done to aggregate signal across similar observations, so if the differences are observed across a sizable region of the graph, the signal would get stronger. The smoothing also depends on the problem, mostly using either graph heat filter or median filter (which is less precise but can be computed much faster). For visualization purposes, we also replaced binary significance values with continuous Z-scores. To estimate those, for each observation difference we found its quantile in the permutation distribution and applied inverse normal density transformation to these quantiles. This yields Z-scores that match the significance levels of the standard normal distribution. In cases where the observations include both positive and negative values, we tested positive values against maximums and negative values against minimums independently.

### Cluster-based expression shifts

Expression differences between conditions for each cell population *C* were determined using the following procedure (more details below the outline):

1. Samples with low number of cells (< 10) for type *C*, as well as genes with low expression (< 1% cells have > 1 UMI) in cell type *C* were filtered out
2. Gene expression counts were summed across all cells within each sample (“pseudo-bulk” measurements), resulting in a count matrix of size ‘*num. samples’* x *‘num. genes’*.
3. Total-count- and log-normalization were applied to this matrix
4. A set of genes, specific for *C* was selected for subsequent analysis (see details below), and the matrix was restricted only to these genes
5. Optionally, dimensionality of data was reduced with principal component analysis (PCA)
6. Pairwise distances 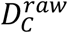 between gene or PCA vectors for all pairs of samples were estimated
7. Expression change was estimated by normalizing 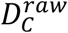 and adjusting it for expected amount of change under permutation of sample labels (see below)
8. Statistical significance of the changes was estimated empirically, based on group randomization

There are several options for selecting genes during step 4. By default, all genes passing the step 1 filtration are selected. However, in some cases it is beneficial to focus on oved-dispersed (OD) or most differentially expressed (DE) genes (see Results). OD genes were estimated using Pagoda2 (https://github.com/kharchenkolab/pagoda2/) implementation, while DE genes were obtained using two-sample Wilcoxon test on log-normalized counts. We did not use more sophisticated approaches for DE, because this test must be repeated for hundreds of permutations (see below), and the packages such as DESeq2 are too slow for such use case. The low quality of the results was partially mitigated as the number of selected genes was fixed *a-priori*, so we did not rely on p-values, provided by the test. The distances between samples 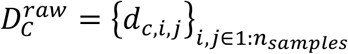, where *n*_*samples*_ is the number of analyzed biological samples, were calculated using either *L*_1_ (Manhattan) or Correlation distance on gene expression or PCA vectors. By default, *L*_1_ distance is used if the number genes / PCs after filtration is less than 30, and Correlation distance is used otherwise.

Next, to normalize the distances between cell types, 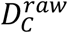 were used to estimate normalized expression shift vectors, with the exact formula varying depending on desired analysis (Fig. 4B).

- 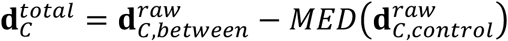: total expression distance, which captures both mean shift of expression and changes in the variance of expression (see Simulation Note)
- 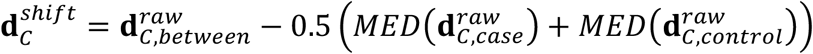: mean expression shifts, which is insensitive to variance changes and mainly captures difference of means
- 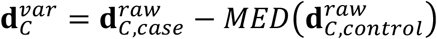: variance distance, which is insensitive to mean shifts and mainly captures difference of variance

Here, *MED* defines median, **d**_*C,between*_ = {*d*_*c,i,j*_}_*i*∈*control,j*∈*case*,_ is a vector of pairwise distances between samples from different conditions, **d**_*C,case*_ = {*d*_*c,i,j*_}_*i*∈*case,j*∈*case,i*≠*j*_ and **d**_*C,control*_ = {*d*_*c,i,j*_}_*i*∈*control,j*∈*control, i*≠*j*_.

Finally, to adjust for other possible random effects and to estimate statistical significance of the results we performed 1000 random permutations of the condition labels for samples within each cell type and estimated the null distribution 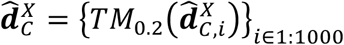, of corresponding 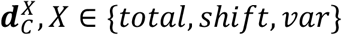. Here, *TM*_0.2_ is the 20% trimmed mean. On this step, only labels of the samples present for cell type *C* were shuffled, so the proportion of samples from each of the conditions was fixed. Next, to obtain the final values, median of this distribution was subtracted from the distances to obtain final values 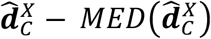 (Fig. 4B). P-values were estimated empirically, as proportion of permutations with distances of the same or greater magnitude as observed: 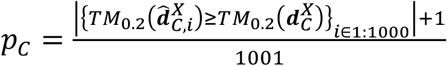 and were adjusted for multiple comparisons using the Benjamini-Hochberg correction. It is important to note that when only top DE genes are used, the DE results depend on the sample labels, and so DE must be re-estimated for each sample label permutation.

### Analysis of sample heterogeneity

To visualize variability of samples we projected distances between samples to 2D space using Multidimensional Scaling (R function *stats::cmdscale*). Different types of distances between samples can be used: it can be the compositional distance D^*coda*^, the cell type-specific expression distance 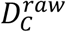, or an overall expression distance D^*raw*^. The first two were described above. The overall expression distances between samples (Fig. 4A,B) were determined as a normalized weighted sum of correlation distances across all cell subpopulations contained in both samples. The base correlation distances *d*^*raw*^ were calculated in the log-normalized expression space, as determined by pagoda2. The weight was equal to the subpopulation proportion, measured as a minimal proportion that the given cell subpopulation represents among the two samples being compared:

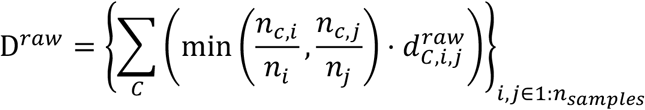

The distances D^*raw*^ or D^*coda*^ are also used for testing separation of samples by user-provided metadata. That is done using Graph Signal Processing framework, by interpreting similarities between sample as a graph adjacency matrix, metadata as a signal on graph, and then estimating whether total variation of this signal on this graph is lower than expected (Shuman et al., 2013). To obtain the adjacency matrix from a distance matrix *D*, the distance matrix *D*^*scaled*-^ is estimated by winsorizing *D* by 5% from each side, and then rescaling it to [0; 1] using min-max scaling. The adjacency matrix is then estimated as *A* = 1 − *D*^*scaled*^. The total variation of the signal ***s*** on graph *A* is estimated as 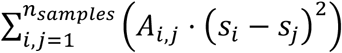 for continuous metadata or as 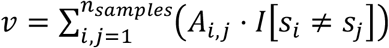 for discrete metadata, where *I*[*c*] is the indicator function. If the metadata has missing values, they are filled with mode for discrete metadata or with median for continuous metadata. Finally, the null distribution of the total variation 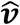 is estimated using the same formulas, by performing 5000 permutations of metadata values across the samples. The significance is estimated as a fraction of randomized cases in which the randomized variation is smaller or equal than the originally observed variation *v*.

### Cluster-free expression shifts

The procedures for estimating the cluster-free expression shifts closely resemble those for the cluster-based version, but are applied to the neighborhoods of individual cells instead of cell types. In addition to the expression matrix, the algorithm requires cell similarity graph as input. In all results presented in the manuscript, the graph was estimated using Conos *buildGraph* method (Barkas et al., 2019), however other methods of constructing a joint graph across collections of samples can also be used. Based on this graph, for each cell *C* the algorithm groups its neighbors by sample, and aggregates molecular counts within each sample, generating a “pseudo-bulk” neighborhood count matrix of size ‘*num. samples’* x *‘num. genes’*. Here, a cell is considered its own neighbor. The samples that had less than 3 cells in the neighborhood of a given cell were omitted from the analysis for this cell. The algorithm requires having at least 2 samples present for each of the conditions, otherwise the cell is excluded from the analysis and the expression shift is set to NA.

From the neighborhood count matrix for the cell *C*, the algorithm estimates pairwise correlation distances between samples,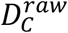. In contrast to the cluster-based version, no gene selection or PCA reduction is available here, and so *L*_1_ distance also cannot be used. Besides that, the expression shift values were estimated using the formulas from the step 7 of the cluster-based expression shifts. The only difference was in permutation of condition labels per sample for estimating the null distribution. Here, each permutation was performed once for the whole dataset, instead of focusing only on samples present in the neighborhood of each individual cells. It resulted in lower precision of these estimates, but enabled much faster computations. The major difference of the cluster-free shifts from the cluster-based version is in the estimation of the p-values (step 8). Like in other cluster-free methods, we applied the correction based on the distribution of the maximal statistics, as described above. Here, smoothing was done using the median filter, and the winsorizing value was set to 0.025. For visualization purposes, the expression shift magnitudes were additionally smoothed over the graph, using the median filter. This allowed them to better match the p-values, which were estimated on these smoothed values.

### Cluster-based differential expression analysis

For the biological results, present in the paper, we used DESeq2 package (Love et al., 2014), using Wald test with parameter *independentFiltering=TRUE*. Then, we filtered genes based on the adjusted p-value threshold of 0.05 (no log2-fold change filtration was done).

### Cluster-free differential expression analysis

Estimation of cluster-free differential expression is very similar to cluster-free expression shifts. The algorithm starts with a cell graph (*Conos buildGraph* was used in this paper) and an expression matrix. Then, for each cell *C*, it splits cells that are adjacent to it by samples and aggregates expression across all adjacent cells within the same sample, thus generating a matrix of size 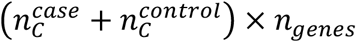. Here 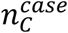 and 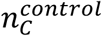 are the number of case and control samples correspondingly, connected to *C*, and *n*_*genes*_ is the number of genes measured in the dataset. If 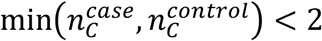, the cell is excluded from the analysis and all Z-scores are set to NA. All samples that have less than 2 cells are excluded from the analysis for *C*. The matrix is total-count normalized, and Z-score for gene *G* is estimated as:

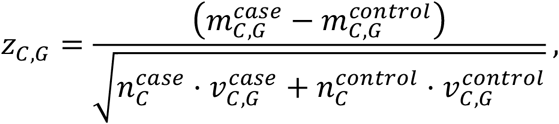

where 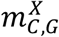 and 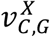 denote mean and variance expression in the condition *X* correspondingly. Z-scores are estimated only for cells that express gene *G* and are set to zero otherwise. Statistical significance is evaluated using the graph-based procedure described earlier, using median smoothing over cell graph with the winsorizing leven of 1%.

To estimate gene programs, we performed clustering of the adjusted z-scores. First, all z-scores < 0.5 were set to 0. Then the clustering was done on top-N genes with the highest sum of absolute z-scores across all cells: 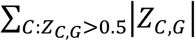. Then, either PAM or Leiden clustering of the z-score matrix for these genes was performed, using cells as features. For PAM, pairwise correlation distances between genes were estimated, and clustering was performed using *cluster* R package for a given number of programs. For Leiden, PCA dimensionality reduction was performed, down to to 100 PCs, using *irlba* package. Then, a k-NN graph was built using N2R package for *k* = 30 and Leiden clustering was run for given resolution and *n. iterations* = 10 using *leidenAlg* R package. Given the gene clustering, program scores were then estimated for each cell as 10% trimmed mean of absolute z-scores across all genes from the given cluster. Finally, genes belonging to each program were ranked by their cosine similarity to the program score.

When estimating global programs we used smoothed adjusted z-scores and Leiden clustering over top-1000 genes with resolution=1. For local programs we used not-smoothed adjusted z-scores and PAM clustering over top-200 genes for the given cell type setting *n. programs* = 4.

### Gene ontology analysis

For Gene Ontology analysis, Cacoa uses clusterProfiler (Wu et al., 2021) functions for Disease Ontology (DO), Gene Ontology (GO) and Gene Set Enrichment Analysis (GSEA). In all cases, Cacoa estimates gene universe as the set of all genes, expressed in at least 5% of cells of the analyzed cell type. For cluster-based analysis, the results presented in the manuscript were performed using GSEA (clusterProfiler wrapper around fgsea (Korotkevich et al., 2016)) with default parameters. For cluster-free analysis we used GO over top-100 genes per gene program for global programs and top-20 genes for local programs.

To analyze ontology enrichment patterns across cell types we visualized heatmaps of enrichment p-values per ontology term (row) per type (column). In such visualization all p-values were re-adjusted across all cell types using Benjamini-Hochberg correction and then converted to -log10-scale. To remove redundant terms we first aggregated terms based on their shared gene content. Specifically, we first estimated individual pathway clusters for each cell type using Jaccard distance on the sets of enriched genes (R functions *hclust* and *cutree* with parameter *h=0.66*). Then, for each pair of pathways (*P1, P2*) we found all cell types that had both *P1* and *P2* enriched and estimated the fraction of cell types that assigned *P1* and *P2* to different clusters: 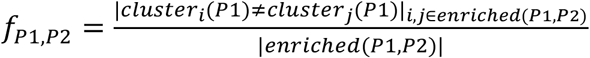. This fraction was used as a distance metric for hierarchical clustering (R functions hclust with parameter *method=“average”* and *cutree* with parameter *h=0.5*). The name of the resulting cluster was selected to have the highest semantic similarity to all cluster members. For that we split the ontology names into words, estimated pairwise Jaccard similarity on words and picked the term with the highest sum similarity. The p-value of the cluster was set to the minimum p-value across all cluster members. We additionally implemented collapsing of ontology heatmaps by cell type patterns (Fig. 5G). First, we trimmed the -log10 p-values by the maximum value of 10. Then we estimated distances between terms using *L1* (Manhattan) distance between the trimmed vectors and performed hierarchical clustering on them (R function hclust, *method=‘complete’*) with the number of clusters being the visualization parameter. Raw log10 p-values (before trimming) were averaged within the cluster to estimate the final pattern, and the name of the cluster was set to the 5 most frequent words across all term names.

### Simulations

To analyze different metrics of expression change we performed simulations with the muscat package (Crowell et al., 2020). We used the Autism dataset to fit the simulation parameters, as it has the largest number of samples per condition. In most analyses, ‘IN-PV’ cell type was used, unless specified otherwise. We simulated data varying different parameters independently to test sensitivity of various expression distance methods to these parameters. For the methods for estimating expression distance, we used: (i) cluster-based expression shifts, (ii) cluster-based expression shifts on top-500 DE genes with PCA reduction to 6 components, (iii) cluster-free expression shifts and (iv) number of differentially expressed genes (using DESeq2 (Love et al., 2014)).

For all analyses, the default parameters were: log2-fold change (LFC)=1.5, fraction of DE genes=0.1, number of cells = 100, number of samples for each condition = 8. Then, we varied each of these parameters independently to test sensitivity for it, keeping the other parameters at their default values. To test for dependency on the number of samples we used values of [3, 5, 7, 9], and [25, 50, 75, 100, 125, 150] for the number of cells per sample. We additionally tested sensitivity of the methods to cell-type specific effects. For that, we compared the expression distance estimates for ‘AST-FB’, ‘AST-PP’, ‘IN-PV’, ‘IN-SST’, ‘L2/3’, ‘L5/6-CC’, ‘Microglia’, ‘Neu-NRGN’ and ‘OPC’ cell types. To evaluate the sensitivity to detect changes in the fraction of DE genes and log2-fold change, we first fixed LFC to 1.0 and varied DE fraction from 0.0 to 0.15 with the step of 0.025. Then, we fixed DE fraction to 0.05 and varied LFC from 1 to 2 with the step of 0.25, as muscat does not allow setting LFC<1.

### scRNA-seq data pre-processing

In pre-processing published datasets, we tried to follow the procedures in the original publications as closely as possible, adding new filtration only in the cases where it was most necessary. For all datasets we removed genes, expressed in less than 10 cells to reduce the matrix size, and removed all mitochondrial genes. Then:

- **Alzheimer**. Cells with fraction of mitochondrial UMIs >0.1 were removed in the original publication. The cells, which contained molecules from multiple donors were removed based on SNPs. We additionally filtered all cells with <500 UMI, and removed doublets using Scrublet (Wolock et al., 2019) on sample expression matrices with threshold of 0.3. We used low-resolution annotation provided by the paper, as the published high-resolution “cell types” were specific to individual samples and were likely driven by batch-effect since no batch-correction algorithms were applied in the paper.
- **Autism**. Cells with fraction of mitochondrial UMIs >0.05 and number of genes <500 were removed in the original publication. We additionally removed doublets using Scrublet on expression matrices with cells grouped by capture batch with threshold of 0.17. The cell annotation was adjusted to merge Neu-NRGN-I and Neu-NRGN-II cell types into Neu-NRGN. This was done because the separation between the two cell types was driven by the number of expressed genes. This separation disappeared when we applied log-transformation to the count data, which was not used in the original publication. When processing the dataset, we noted that while only two brain regions were reported in the manuscript, the sample names encoded four different regions, which were clearly separated by gene expression (see the Results). So, for testing separation of samples by metadata we extracted high-resolution region names from the cell ids, assuming the encoding was done as *‘individual_region’* (*e.g. ‘6033_BA24’*).
- **Epilepsy**. All the QC filtration was already performed in the original publications. There, cells were filtered based on minimal number of UMIs (individual thresholds per sample) and fraction of mitochondrial UMIs >0.08. And doublets were removed using Scrublet and expression of mutually exclusive markers. We therefore used the filtered data and the highest-resolution annotation provided in the original manuscript.
- **MS**. Cells with less than 500 genes or 1000 UMIs were removed in the original publication. We additionally removed cells with the mitochondria fraction >0.1, and filtered doublets using Scrublet with threshold of 0.25. The authors separated cells specific to Control or MS for ‘L2-L3 EN’ and ‘OL’ subtypes. For an unbiased analysis we modified the annotation to remove that separation.
- **PF**. All QC filtration was already performed in the publication. We took all the cells reported in the publication from the “Control” and “IPF” conditions.
- **SCC**. We followed the filtration procedures described in the original publications: the cells with mitochondrial fraction >0.1, as well as ones annotated as ‘Multiplet’ were removed. We also removed the cell cluster annotated as ‘Keratinocyte’, containing Keratinocytes removed in the original publication, even though this introduces bias in the downstream analysis. In the original publication, the authors have “removed a small number of cells coming from tumor samples that clustered with normal cells and vice versa” (Ji et al., 2020). Such filtration artificially increases significance of some of the differentially expressed genes, increases expression shift magnitudes, and distorts cell counts in compositional analysis. We maintained this filtering step purely for the consistency with the publication. Finally, we removed three samples (*P4_Normal, P8_Normal* and *P3_Tumor*) with very low number of cells after filtration (22, 349 and 419 cells respectively).

On the filtered counts, Pagoda2 (https://github.com/kharchenkolab/pagoda2/) objects were created using parameters *trim=10* and *log.scale=TRUE*. Gene variance was adjusted using *gam.k=10*, and then PCA reduction was estimated using top 3000 over-dispersed genes and 100 first principal components. These Pagoda2 objects were used to perform alignment with Conos (Barkas et al., 2019), using graph parameters *k=20* (*k=40* for EP), *k.self=5, space=‘PCA’, ncomps=30, n.odgenes=2000, matching.method=‘mNN’* and *metric=‘angular’*. The graph embedding was estimated using UMAP with default parameters, with the only exception of the SCC dataset, where we set *spread=10*.

### Stability of Differential Expression Analysis

To assess stability of the DE results for different methods and cell types, we employed a leave-one-out (LOO) resampling procedure: multiple runs of DE analysis were run iteratively, leaving out one sample at a time. For each iteration, we obtained DE genes (defined by top N genes or a p-value cutoff), and then calculated the Jaccard coefficients for each pair of the resulting DE gene sets. We then summarized the Jaccard coefficients per cell type as a mean value across pairwise comparisons. The mean value of Jaccard coefficients across cell types was used as a characteristic of a particular DE method on a specific dataset. The stability per gene was estimated as a mean p-value rank of a gene across all LOO DE results.

Looking to increase DE stability, we also devised a modified DE procedure, relying on the LOO sampling internally (“LOO:DESeq2”). Specifically, the procedure wrapped an existing DE method (DESeq2, edgeR, limma-voom, t-test, and Wilcoxon test), by running LOO procedure within the provided set of samples, and returning a DE list sorted based on the mean p-value rank of each gene across all LOO iterations. Such wrapped methods were prefixed with “LOO” (i.e. “LOO:DESeq2”). The stability of the wrapped method ((“LOO:DESeq2”) was evaluated in the same way as others, that is by putting it through an outside LOO procedure to determine its overall stability characteristics.

### Identification of latent covariates optimizing DE stability with Genetic Algorithm

We hypothesized that the instability of the DE results may be associated with some unobserved study co-variate. We therefore designed an approach to identify a sample-level latent covariate that would optimize the stability of the DE results.

For each cell type, we optimized a latent covariate (vector of continuous values of the same length as the number of samples) using a genetic algorithm (GA). The fitness function (objective function) of the GA was taken to be the minimum Jaccard index across 100, 200, 300, 400, and 500 top genes after the LOO procedure. Each DE calculation used DESeq-Wald method with the latent factor passed as a sample covariate. The population size in the GA was set to 90. The 30 best individuals were taken as parents to form the new generation. A new individual was produced as a recombination of two random parents with noise. The number of epochs was set to 60.

## Code availability

The methods described in this manuscript are implemented in the Cacoa R package (https://github.com/kharchenkolab/cacoa/). The code to reproduce all analyses from the manuscript is available on GitHub (https://github.com/kharchenkolab/cacoaAnalysis) with the workflow [<u>ref</u>] web-site containing the compiled R notebooks and individual dataset reports (https://kharchenkolab.github.io/cacoaAnalysis/)

### Datasets

1. <u>Alzheimer</u>: Grubman, A., Chew, G., Ouyang, J. F., Sun, G., Choo, X. Y., McLean, C., …Polo, J. M. (2019). A single-cell atlas of entorhinal cortex from individuals with Alzheimer’s disease reveals cell-type-specific gene expression regulation - Nature Neuroscience. Nat. Neurosci., 22, 2087–2097. doi: 10.1038/s41593-019-0539-4
2. <u>Autism</u>: Velmeshev, D., Schirmer, L., Jung, D., Haeussler, M., Perez, Y., Mayer, S., …Kriegstein, A. R. (2019). Single-cell genomics identifies cell type–specific molecular changes in autism. Science.
3. <u>Epilepsy</u>: Pfisterer, U., Petukhov, V., Demharter, S., Meichsner, J., Thompson, J. J., Batiuk, M. Y., …Khodosevich, K. (2020). Identification of epilepsy-associated neuronal subtypes and gene expression underlying epileptogenesis - Nature Communications. Nat. Commun., 11(5038), 1– 19. doi: 10.1038/s41467-020-18752-7
4. <u>MS</u>: Schrimer et al. “Neuronal vulnerability and multilineage diversity in multiple sclerosis”, Nature 2019 Sep;573(7772):75-82. doi: 10.1038/s41586-019-1404-z. ACCESSION ID
5. <u>PF</u>: Habermann, A. C., Gutierrez, A. J., Bui, L. T., Yahn, S. L., Winters, N. I., Calvi, C. L., …Kropski, J. A. (2020). Single-cell RNA sequencing reveals profibrotic roles of distinct epithelial and mesenchymal lineages in pulmonary fibrosis. Sci. Adv. Retrieved from https://www.science.org/doi/10.1126/sciadv.aba1972
6. <u>SCC</u>: Ji, A. L., Rubin, A. J., Thrane, K., Jiang, S., Reynolds, D. L., Meyers, R. M., …Khavari, P. A. (2020). Multimodal Analysis of Composition and Spatial Architecture in Human Squamous Cell Carcinoma. Cell, 182(2), 497–514.e22. doi: 10.1016/j.cell.2020.05.039

