## Supplemental Figures for "Case-control analysis of single-cell RNA-seq studies"

Supplementary figures 1-5.

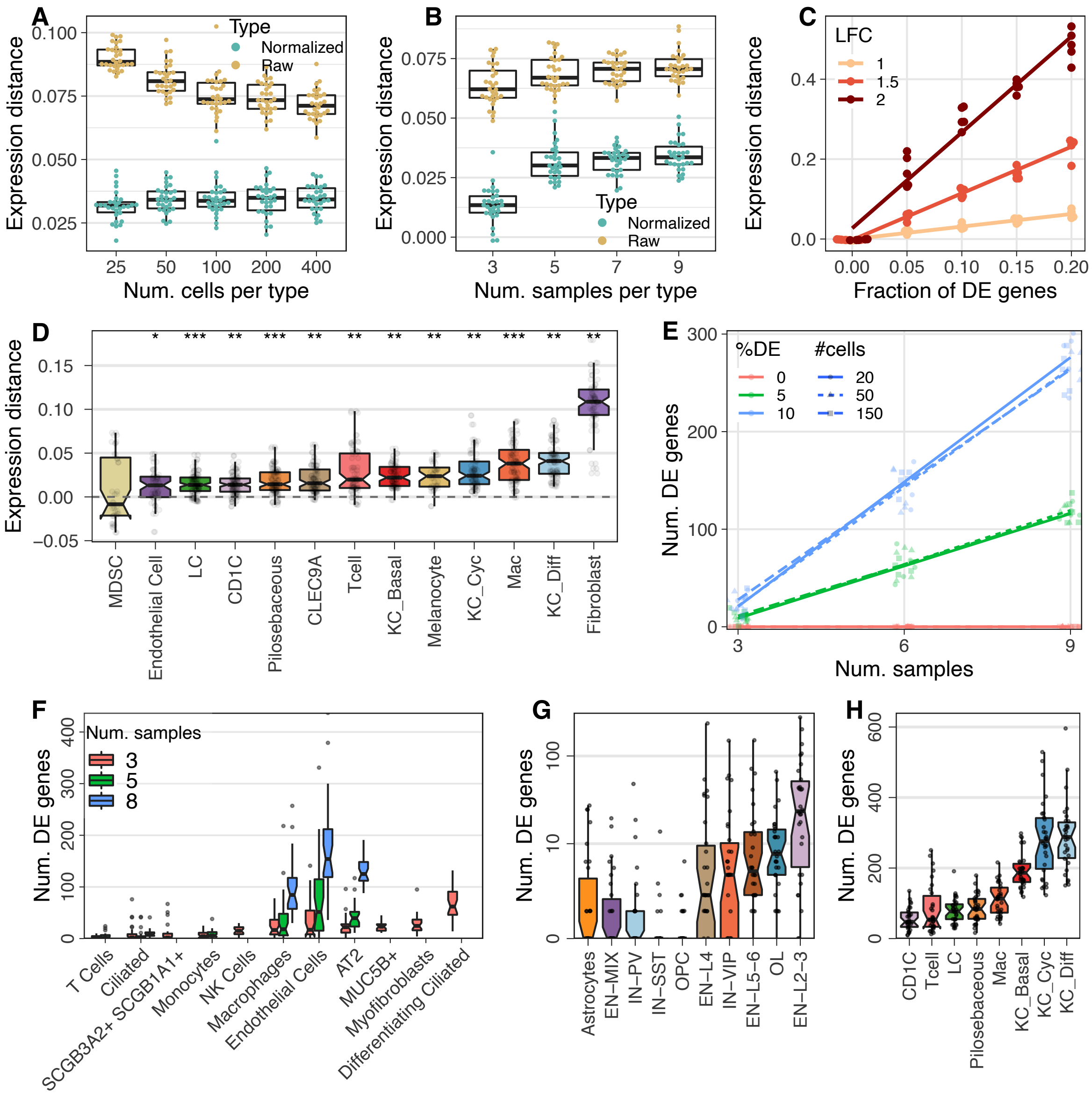


**Supp. Figure 1. Cluster-based analysis of expression change magnitudes.**
(A-B) Sensitivity of the expression distance (y-axis) before normalization (yellow) and after normalization ("expression shift", green) to different covariates (x-axis): number of cells (A) and number of samples (B). The data was simulated using muscat, and each dot corresponds to the median distance across all pairs of samples for one simulation. The normalization removes dependency on the number of cells, and the number of samples does not influence it except for extremely low values.
(C) Sensitivity of the expression shift metric (y-axis) to changes in log2-fold change (color) and fraction of differentially expressed genes (x-axis). Each point is a median distance for one simulation, with lines showing a linear regression fit.
(D) Visualization of expression shifts for the SCC dataset in the same format as Figure 4D.
(E) Dependency of the number of significant DE genes (using DESeq2 with BH adjustment, y-axis) on the number of samples (x-axis) for the different numbers of cells (line type) and the fractions of simulated DE genes (color). The plot is in the same format as Fig. 4H, but the number of cells is controlled for by subsampling. The number of samples still strongly affects the results.
(F-H) Number of significant DE genes (y-axis) for different cell types (x-axis) after subsampling of samples for PF (F), MS (G) and SCC (H). Both biological samples and cells were down-sampled to control for these covariates. Each dot corresponds to one down-sampling result. (F) shows a trade-off between the number of biological samples used for DE estimation (color) and the number of cell types having the required number of samples. Increasing the number of samples available for down-sampling enables comparison of more cell types, but shows higher variability and lower sensitivity.

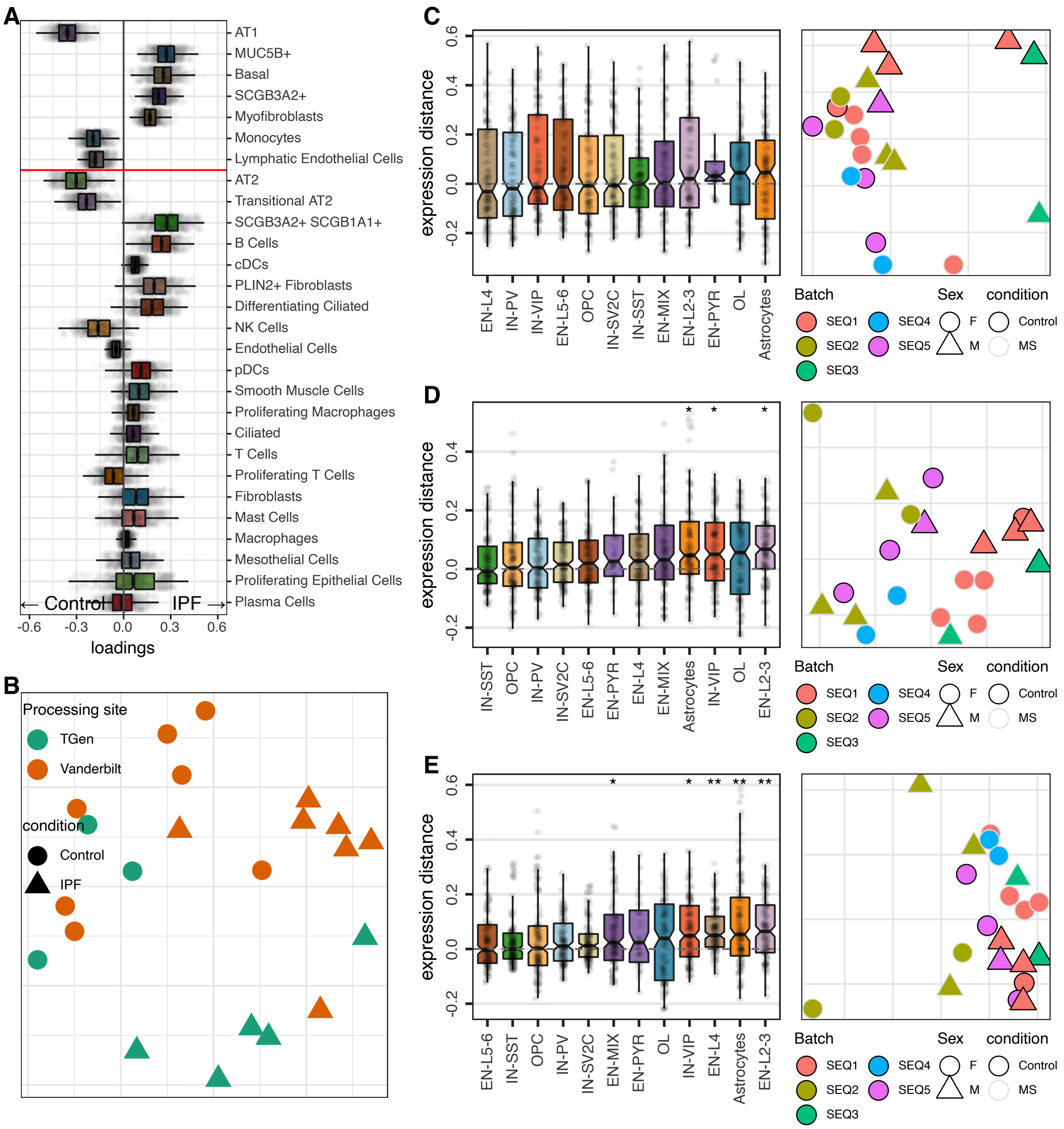


**Supp. Figure 2. Analysis of sample heterogeneity.**
(A) Cluster-based composition analysis applied to the PF dataset using only data from the Vanderbilt processing site. In contrast to the full dataset results (Fig. 1H), it shows Fibroblast subtypes over-represented in IPF.
(B) MDS embedding of the CoDA space for the PF dataset. The format is the same as for Fig. 4E.
(C-E) Visualization of expression shifts (left) estimated on over-dispersed (OD) genes and the corresponding sample MDS embedding (right) for the MS dataset. The format is the same as on Fig. 4D,E. MDS plots are colored by Batch, point shape corresponding to Sex, with Control samples having black outlines. (C) shows the distances on top-100 OD genes and it has clear separation by Sex. (D) shows distances for top-500 OD genes with sex-related genes filtered out (see Methods). Here, Sex differences were removed, while Batch differences appear as the main driver of differences. Such filtration of genes increased sensitivity of the expression shifts (left). (E) distances filtering out both Batch- and Sex-related genes. While there is still visible separation by Batch, it has the highest sensitivity of the three.

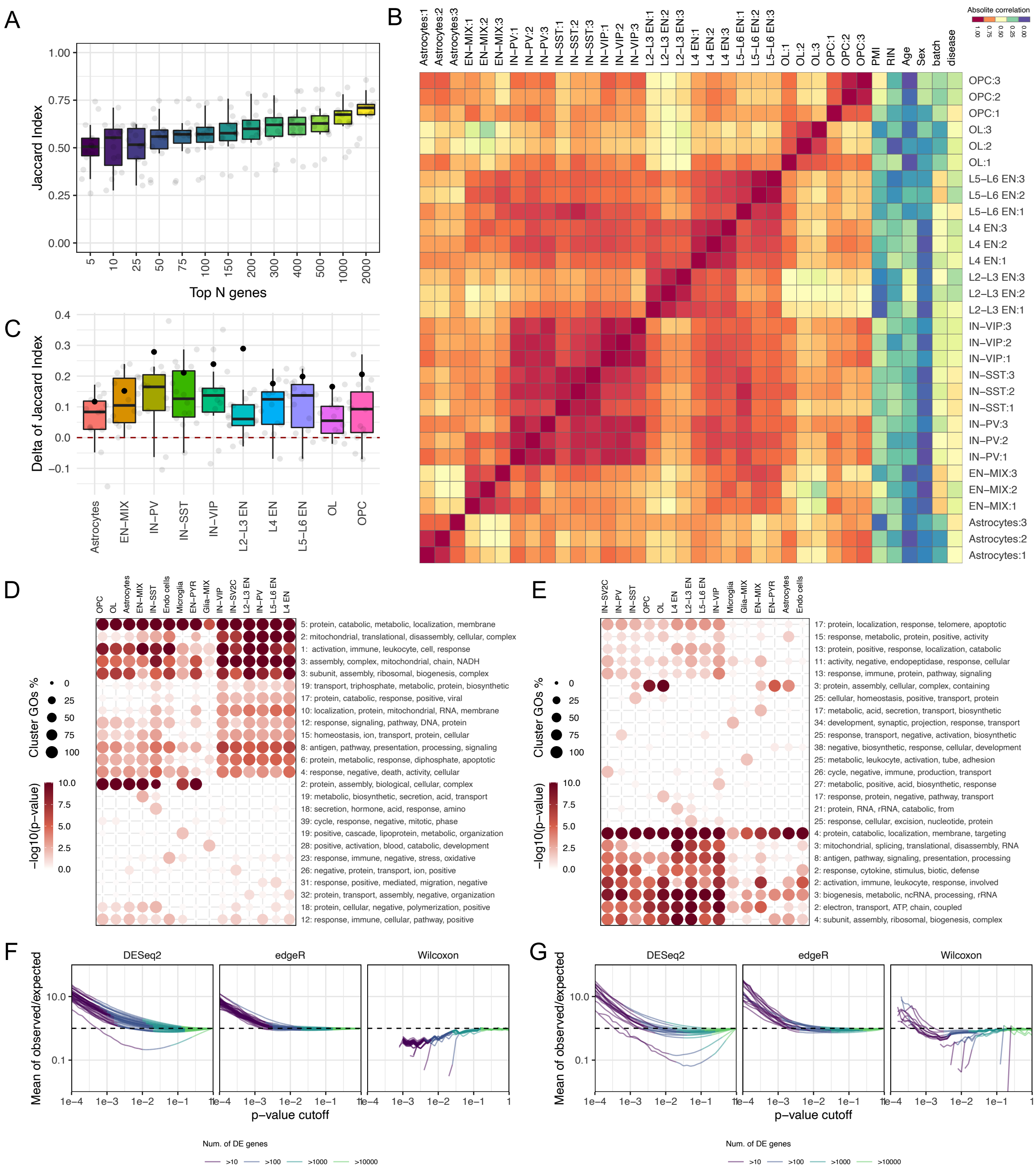


**Supp. Figure 3. Differential expression analysis.**
(A) Jaccard Index between sets of top-N DE genes after leave-one-out procedure (y-axis) is shown for the MS dataset, as a function of N (x-axis).
(B) Absolute value of correlation of latent factors optimizing DE gene set stability for different cell types (rows) is shown with respect to each other and to covariate factors tracked in the study (columns).
(C) Improvement in the top-N DE gene set stability (N=100) is shown for all other cell types (gray dots) for the latent factors optimized for different target cell types (x-axis). The improvement of the target cell type itself is shown with a black dot. The boxplot shapes consider only gray dots (*i.e.* exclude the target cell type). The improvement for all non-target cell types is statistically significant (Student t test P<0.001 after multiple hypothesis testing corrections).
(D-E) Overview of top GSEA sets for the DE genes determined for different cell types on the original dataset (E), and with inclusion of a stabilizing latent factor (D) optimized for the IN-PV cell type. The format follows Fig. 5G.
(F-G) Similar to the Fig. 5F of the main manuscript, the observed/expected number of DE genes is shown under randomized sample assignment to groups for Epilepsy dataset (F), MS dataset (F).

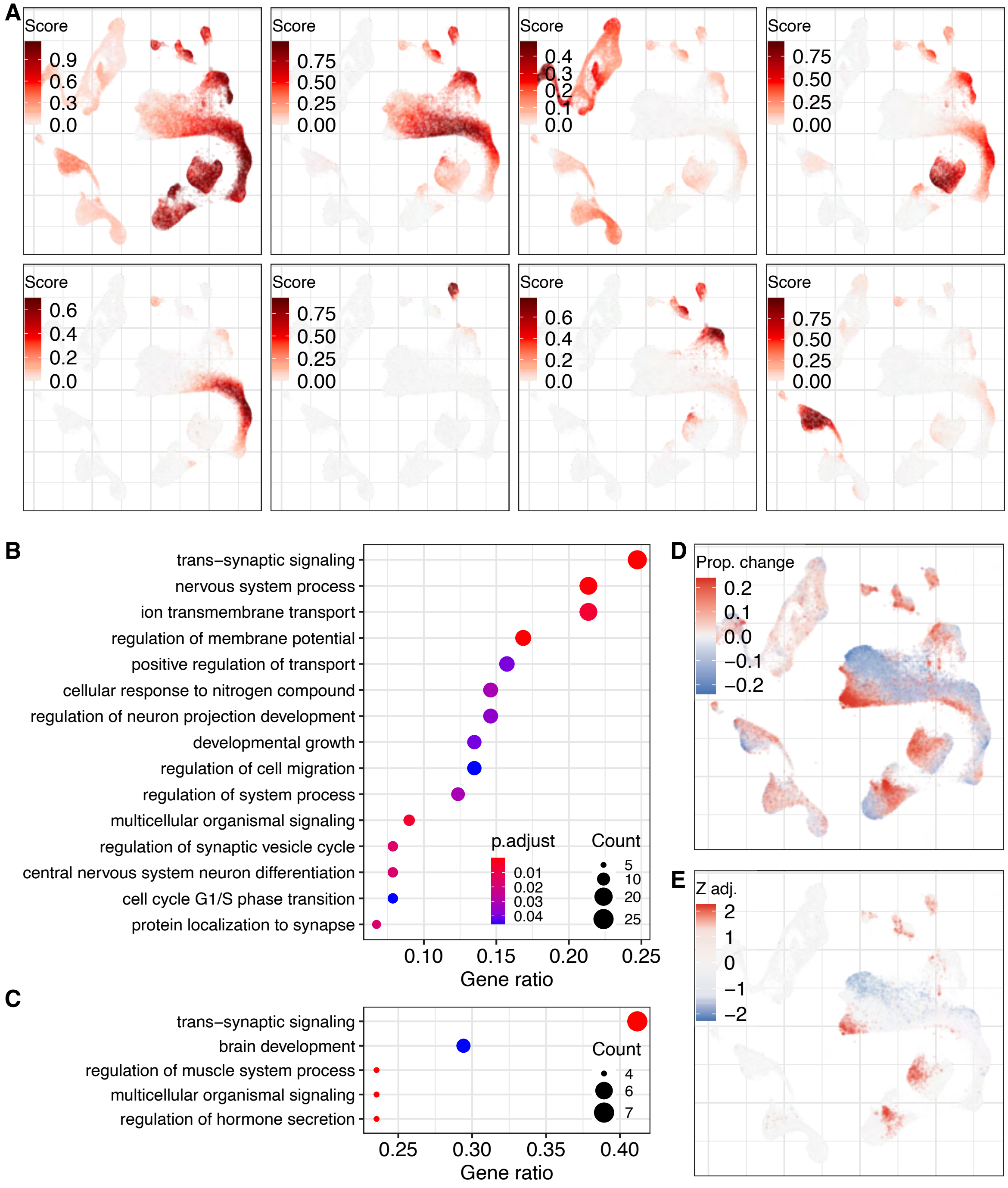


**Supp. Figure 4. Cluster-free gene expression analysis.**
(A) Visualization of the eight most pronounced global gene expression programs identified by Leiden clustering of genes, shown in the same way as Fig. 6K.
(B-C) GO enrichment dotplots, showing clustered ontology term significance (same format as Fig. 7E,F) for the Excitatory neuron gene program from Fig. 6K (B) and the L2_Cux2_Lamp5 local programs from Fig. 6P (C).
(D-E) Cluster-free graph-based compositional changes of the Epilepsy dataset, with the magnitude of change shown in (D) and statistical significance shown in (E). The format is similar to Fig. 3F,H.

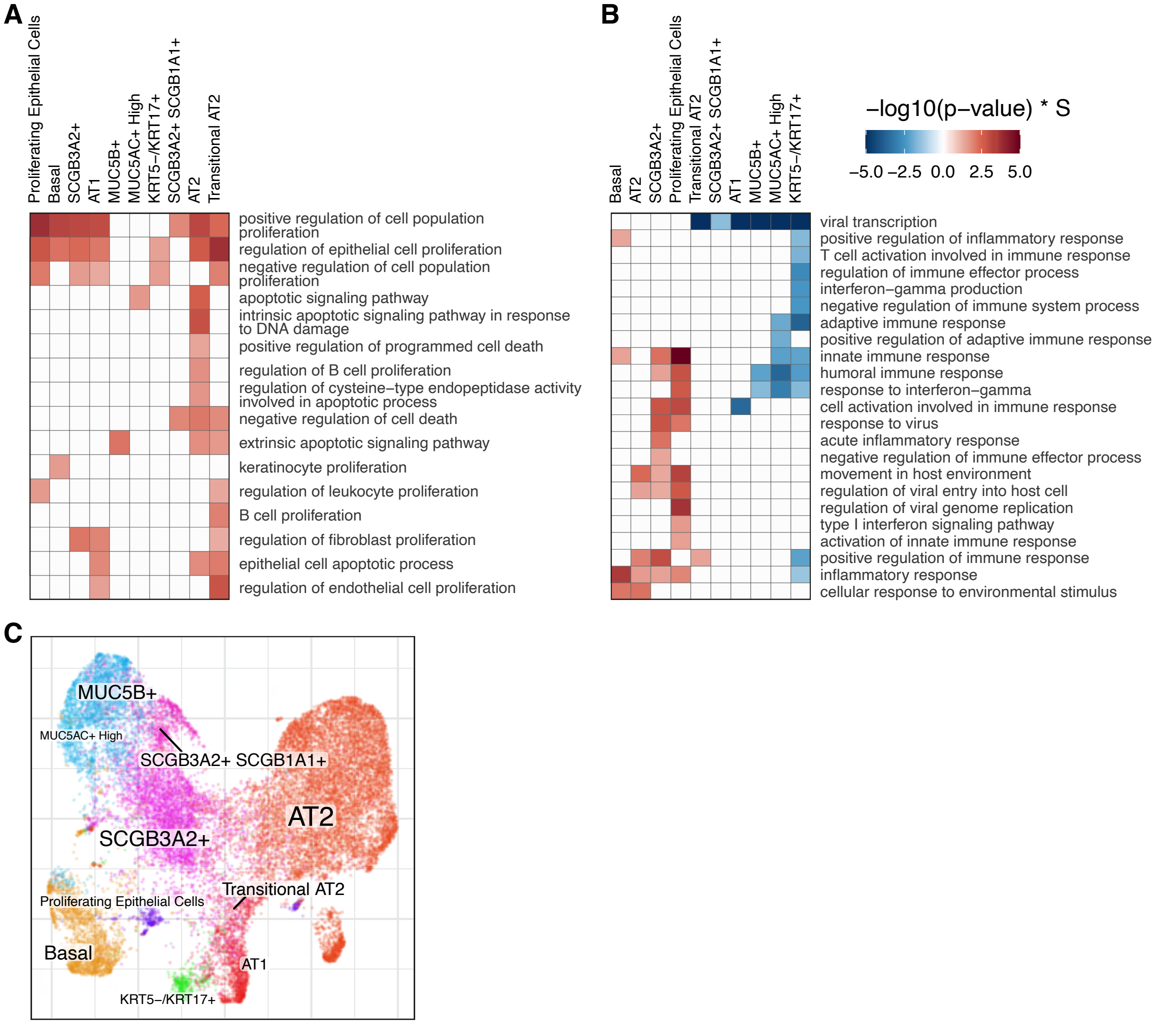


**Supp. Figure 5. Functional interpretation.**
(A-B) Gene Ontology heatmaps in the same format as Fig. 7A. (A) Cell death and proliferation-related ontologies across Epithelial cell types. (B) Virus and immune response ontologies across Epithelial cell types.
(C) Annotated UMAP embedding of Epithelial cell types for the PF dataset.
